# A membrane integral methyltransferase catalysing N-terminal histidine methylation of lytic polysaccharide monooxygenases

**DOI:** 10.1101/2022.10.03.510680

**Authors:** Tanveer S. Batth, Jonas L. Simonsen, Cristina Hernández-Rollán, Søren Brander, Jens Preben Morth, Katja S. Johansen, Morten H. H. Nørholm, Jakob B. Hoof, Jesper V. Olsen

## Abstract

Lytic polysaccharide monooxygenases (LPMOs) are oxidative enzymes that help break down lignocellulose, making them highly attractive for improving biomass utilization in biotechnological purposes. The catalytically essential N-terminal histidine (His1) of LPMOs is post-translationally modified by methylation in filamentous fungi to protect them from auto-oxidative inactivation, however, the responsible methyltransferase enzyme is unknown. Using mass-spectrometry-based quantitative proteomics in combination with systematic CRISPR/Cas9 knockout screening in *Aspergillus nidulans*, we identified the N-terminal histidine methyltransferase (NHMT) encoded by the *gene* AN4663. Targeted proteomics confirmed that NHMT was solely responsible for His1 methylation of LPMOs. NHMT is predicted to encode a unique seven-transmembrane segment anchoring a soluble methyltransferase domain. Co-localization studies showed endoplasmic reticulum residence of NHMT and co-expression in the industrial production yeast *Komagataella phaffii* with LPMOs resulted in His1 methylation of the LPMOs. This demonstrates the biotechnological potential of recombinant production of proteins and peptides harbouring this unique post-translational modification.

## INTRODUCTION

Protein methylation is a post-translational modification (PTM) widely utilized to regulate DNA transcription via the modification of histones at lysine and arginine residues in eukaryotic organisms, constituting the primary component of the so-called “histone code”(*1*). Histidine methylation of proteins was recently described in mammalian cells targeting highly abundant actin proteins and its primary function has been suggested to moderately regulate actin filament(*2*). More recently, the human methyltransferases METTL9 and METTL18 have demonstrated histidine methylation activity. METTL9 was shown to mediate ubiquitous histidine methylation in mammalian proteomes(*3*), whereas METTL18 was shown to be a histidine-specific methyltransferase that specifically targets 60S ribosomal protein L3 and affects ribosome biogenesis and function(*4*). Histidine N-methyltransferase activity has developed on both SET(*2*) and seven-β-strand domain containing(*4*) class of methyltransferases.

The histidine methylation(*5, 6*) found on fungal lytic polysaccharide monooxygenases (LPMOs) is remarkable due to their N-terminal histidine specificity(*7*). LPMOs are a class of oxidative enzymes found widely in nature. They have broad substrate specificity towards complex polysaccharides, including lignocellulose(*8, 9*). Cellulose-specific LPMOs are used commercially for the conversion of plant biomass to biofuels(*10*). LPMOs have also been implicated in several fungal, viral and bacterial mediated infections that can be lethal(*11–13*). However, N-terminal histidine methylation is only observed on LPMOs expressed by filamentous fungi. A defining feature of all LPMOs is the N-terminal histidine, which is crucial for copper binding and enzyme activity. It is this N-terminal histidine residue that is methylated on its imidazole nitrogen (τ-methylation of Nε2, Figure 1A). This PTM prevents protonation of the His1 side chain, and it has been demonstrated that it protects the critical active site of the enzyme from oxidative damage(*14*). Although X-ray crystallography has shown methylation of the N-terminal histidine in several fungal LPMOs (*8, 15*), the methyltransferase responsible for this modification remains unknown.

**Figure 1.**
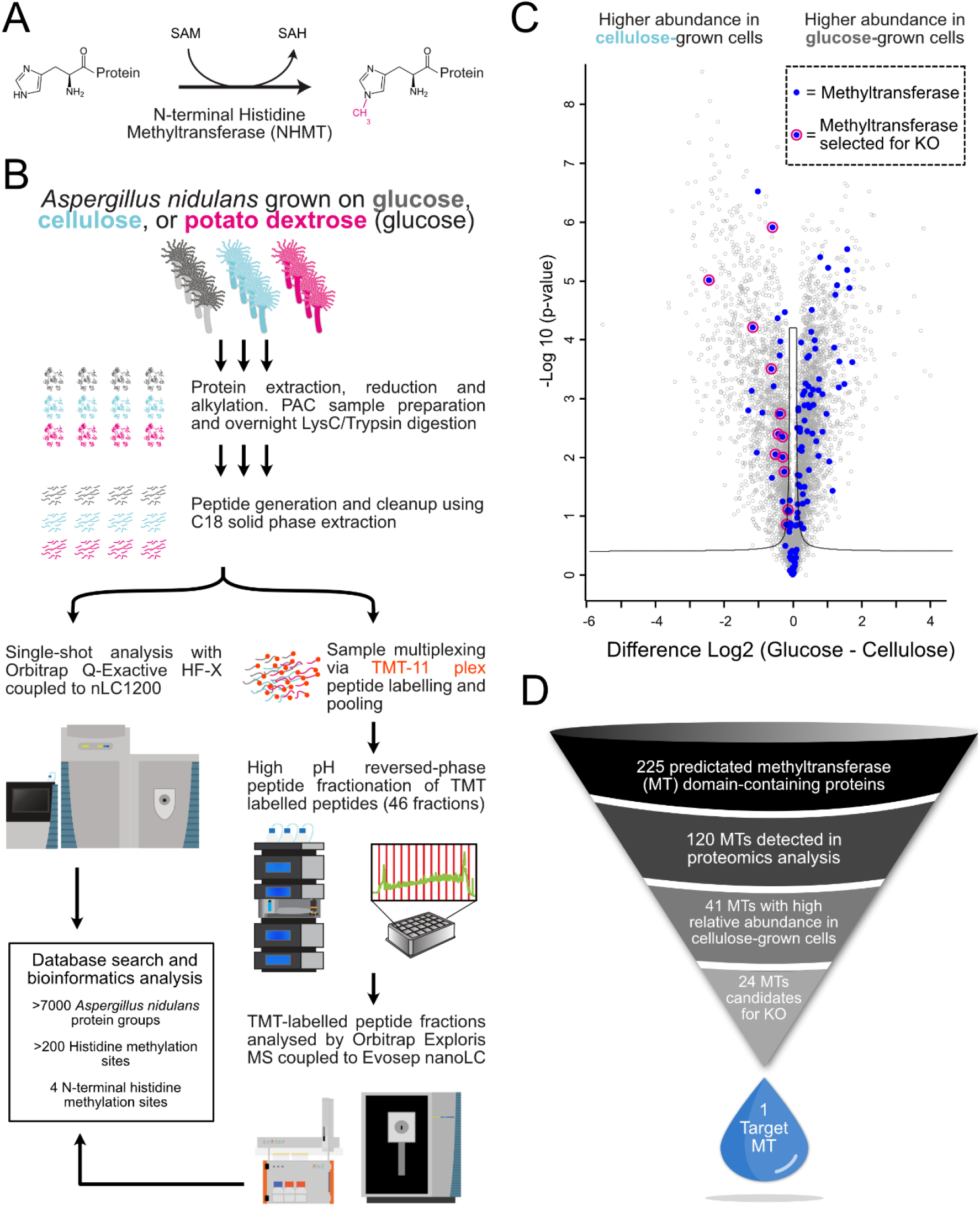
Illustration of the workflow in the NHMT identification. **A)** Mechanism of N-terminal histidine methylation via the transfer of a methyl group from S-adenosylmethionine (SAM). **B)** *A. nidulans* cells were grown on different *carbon sources* followed by protein extraction and LysC/trypsin digestion. The peptides generated were analysed using an Orbitrap-based mass spectrometer. **C)** Volcano plot showing the protein abundance differences (from TMT experiment) of cells grown in medium with cellulose versus cells grown in a medium with glucose. All identified methyltransferases (blue dots) and methyltransferases (blue dots with pink outline) selected for knockout from this dataset are illustrated. **D)** Graphical illustration of the process leading to the knockout(KO) candidates followed by the identification of the NHMT.

In this study, we used state-of-the-art quantitative proteomics in combination with a CRISPR/Cas9-based gene knockout strategy to identify and characterize a previously undescribed N-terminal histidine methyltransferase (NHMT) in the filamentous fungi *Aspergillus nidulans*. We demonstrate that this enzyme is solely responsible for the unique His1 methylation of LPMOs in *A. nidulans*. The methyltransferase is predicted to contain a novel seven helical transmembrane domain and a soluble catalytic domain.

## RESULTS

### Shortlisting N-terminal histidine methylation candidates using bioinformatics and a quantitative proteomics screen

To identify and shortlist NHMT candidates, a large-scale quantitative proteomics screen was performed to identify differentially expressed cellular proteins of cells grown on different carbon sources (Figure 1B). This strategy was based on the hypothesis that NHMT candidates are co-expressed with methylated LPMO substrates, specifically when grown under conditions where the LPMO is needed, such as when *A. nidulans* is grown with cellulose as an additional carbon source.

Differential protein expression analysis of cells grown with the different carbon sources was accomplished through a combination of single-shot label-free quantification (LFQ) and tandem mass tag (TMT)-labelled multiplex in-depth quantitative mass-spectrometry. Additionally, we performed deep proteome sequencing for *A. nidulans*, identifying almost 7000 protein groups and generating the largest dataset of filamentous fungal proteins to date (Supplementary Table S1).

Over 200 histidine methylation sites were identified in the dataset, four of which were localized to N-terminal histidines on four different proteins (Supplementary Table S2). All of the four N-terminally histidine methylated proteins with UniProt identifiers: C8V530, Q5B1W7, Q5AU55, and Q5B428, are predicted LPMOs or LPMO-like proteins of the auxiliary activity family type based on automatic fold recognition and structure prediction (*16*). C8V530, gene ID AN10419, is classified as a member of Auxiliary Activity Family 9 (AA9 LPMO) of the CAZy class of glycoside hydrolases by InterPro (*17*) and Pfam(*18*) (Table S1). Q5B1W7 (AN5463) plays a crucial role in starch degradation (*19*), and similarly contains a predicted glycoside hydrolase domain followed by a predicted carbohydrate-binding module by Pfam (Table S2). Q5AU55 (AN4702) is structurally predicted to be most similar to an AA11-type LPMO from *A. oryzae*. Lastly, Q5B428 (AN8175) is highly homologous to an LPMO-like protein from *Laetisaria arvalis* (LaX325) but contains a predicted transmembrane anchoring sequence at the C-terminal of the protein. All four of these proteins contain a secretion signal peptide prior to the N-terminal histidine of the processed proteins.

To identify NHMT candidates, relative protein abundances in cells grown on glucose-containing media (minimal media with glucose or potato dextrose) were compared with those in cells grown with the addition of cellulose. More specifically, using volcano plot analysis of the log2-fold changes and t-test based statistics; we focused on putative methyltransferases that exhibited different abundances when grown on different media (Figure 1C). To identify all putative methyltransferases in *A. nidulans*, bioinformatic analysis was performed using a combination of Interpro(*17*), Pfam(*18*), PROSITE(*20*) and gene ontology annotations. From this, a total of 225 predicted putative methyltransferase genes were annotated from the *A. nidulans* FASTA protein sequence database (Supplementary Table S3). We ultimately identified 120 of the 225 predicted methyltransferases in the *A. nidulans* proteome analysis, 41 of which were found to have statistically significantly higher abundance when *A. nidulans* was grown on cellulose as the primary carbon source in the growth media compared to glucose or potato dextrose (Figure 1C, Supplementary Table S4). Seven of the 41 methyltransferases were eliminated due to their prediction (based on combined annotations) as methyltransferases with oxygen as the acceptor atom instead of nitrogen. Additional methyltransferases were eliminated due to the existence of homologs in *Saccharomyces cerevisiae* or *Komagataella phaffii* as these organisms are incapable of performing the modification. Moreover, phylogenetic analysis revealed that 19 of the remaining methyltransferases appeared to be unique among fungi known to encode for LPMOs (Supplementary Table S4). To identify the hitherto unknown NHMT enzyme responsible for N-terminal methylation of LPMOs in filamentous fungi, systematic CRISPR/Cas9-based gene knockout screening was carried out on the 19 candidates in *A. nidulans* (Figure 1D). Another five manually curated methyltransferases that lacked homologs in *Saccharomyces cerevisiae* or *Komagataella phaffii* were included. Of the 24 candidates, 22 were successfully knocked out and assayed for NHMT activity.

### CRISPR/Cas9 knockout screening of 22 methyltransferase candidate genes coupled with targeted MS analysis of LPMO methylation status

To precisely and efficiently knock out each of the 22 genes by CRISPR/Cas9, we designed guide RNAs for each candidate and used a Cas9-expressing strain deficient in the error-prone non-homologous end-joining DNA repair, ensuring high-fidelity genome editing through homologous recombination(*21, 22*). To analyse the effect of the individual knockouts on N-terminal histidine methylation, we developed a targeted proteomics assay based on parallel reaction monitoring (PRM) to specifically monitor and quantify the native N-terminally histidine methylated *A. nidulans* peptide sequences identified in the proteome analysis. Of the four N-terminal histidine methylated protein sequences detected in the large-scale proteomics screening, we ultimately selected the most abundant His1 methylated peptide corresponding to ([meth]HTVIVYPGYR) from the uncharacterized *A. nidulans* LPMO-like protein Q5B428 (encoded by the gene AN4702). We prioritized this protein because its N-terminal histidine methylated peptide was reproducibly detected in all proteomics experiments without the need for extensive peptide fractionation and readily detected in rapid single-shot PRM-MS analysis (Figure 2A). We also targeted the predicted unmethylated peptide and required the observation of this to constitute a hit in individual knockout strains (Figure 2A). This quantitative PRM analysis revealed that 21 of the 22 methyltransferase knockouts did not affect the methylation state (Figure 2A, Figure S1, Supplementary Table S5). However, the knockout of the gene encoding for AN4663 (corresponding to the UniProt identifier Q5B467) abolished N-terminal histidine methylation on the monitored methylated peptide and conversely, the unmethylated form was for the first time observed (Figure 2B, C and D).

**Figure 2.**
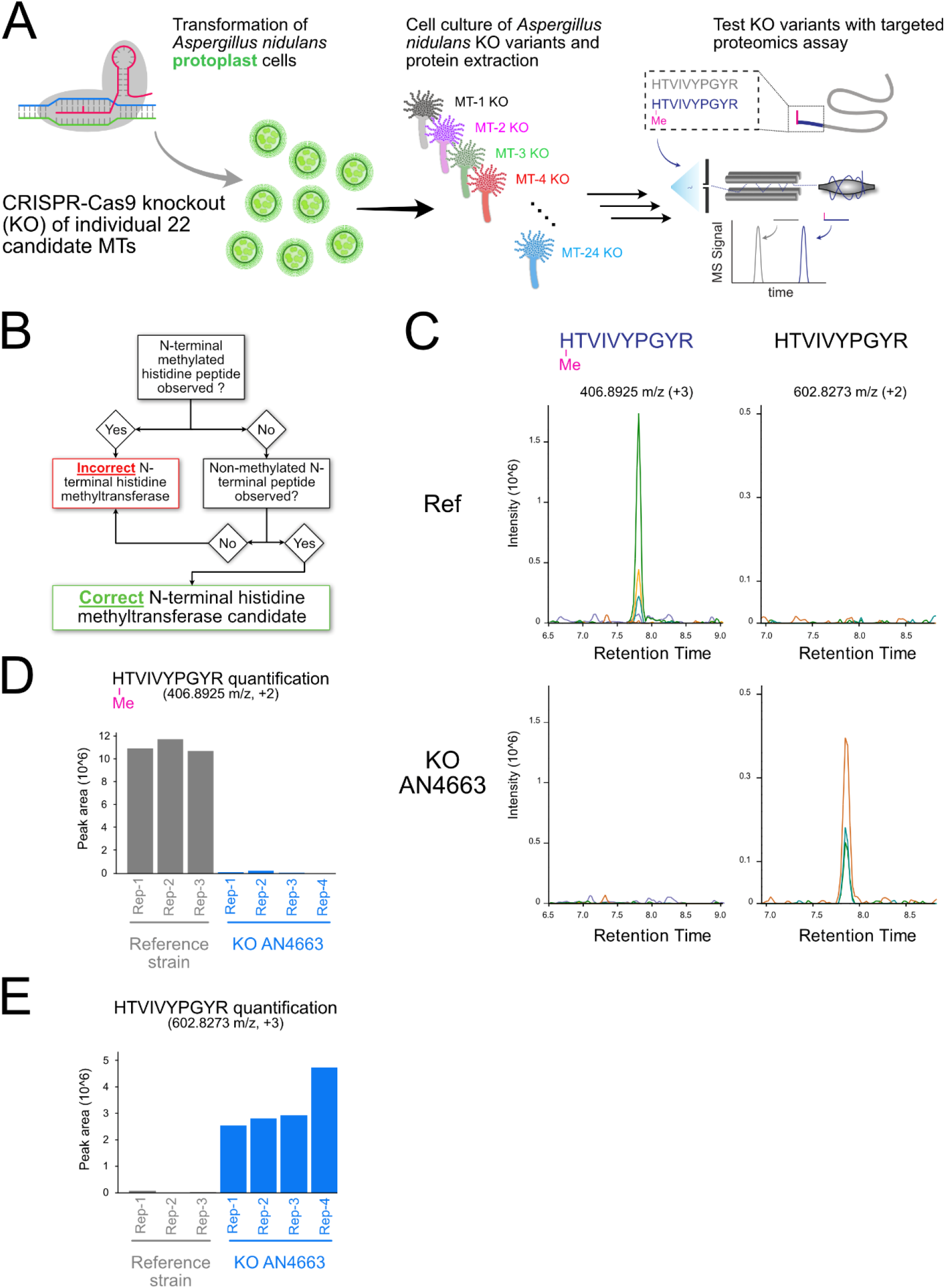
CRISPR/Cas9 knockout and proteomics strategy for identification of the NHMT. **A)** Schematic representation of the CRISPR/Cas9 knockout library generation and subsequent targeted proteomics assay for identification of the NHMT. **B)** Workflow describing the process required to identify the novel NHMT target from the PRM assay. **C)** PRM chromatogram of N-terminal histidine methylated and unmethylated HTVIVYPGYR peptide in reference strain (top) and AN4663 knockout strain (bottom). **D)** The relative quantification of the N-terminal methylation histidine peptide ([meth]HTVIVPGYR) for the reference strain, the knockout candidate AN4663. **E)** Relative quantification as in panel D of the unmethylated counterpart peptide HTVIVYPGYR.

### AN4663 is the specific N-terminal histidine methyltransferase encoding gene in *A. nidulans*

In UniProt, AN4663 has been classified as a spermidine synthase (SpdS) based on automated annotation. *In silico* structural analysis of the 558-amino-acid-long AN4663 protein using PSIPRED(*23*) and the transmembrane protein topology prediction tool using MEMSAT(*24*) revealed an predicted domain architecture with a N-terminal seven-transmembrane (TM) domain up to amino acid position 215, followed by a soluble methyltransferase-active domain at the C-terminal (Figure 3A and Supplementary Figure S2). This C-terminal domain in AN4663 is structurally similar to the SpdS-like domain in the human methyltransferase (MTase)-like protein 13 (METTL13), albeit with low amino acid homology. METTL13 has been demonstrated to be solely responsible for the stoichiometric tri-methylation of the N terminus of eukaryotic elongation factor 1 alpha(*25*). A sequence alignment to METTL13 identified a conserved glutamate at position 340 which is required for the binding of the co-substrate S-adenosylmethionine (SAM) but not the catalytic activity of methyltransferases(*26*). Experimental evidence for the correct identification of NHMT was obtained by site-directed mutagenesis of this glutamate to an alanine (E340A) in *A. nidulans*. This resulted in the complete loss of N-terminal histidine methylation in the His1 peptide monitored, similar to the complete knockout of AN4663 (Figure 3B, C, and D, Supplementary Table S5). Collectively, these data confirm that AN4663, henceforth denoted as *nhmT* when referring to the gene locus, encodes the specific and sole NHMT enzyme in *A. nidulans* responsible for N-terminal histidine methylation of co-expressed LPMOs.

**Figure 3.**
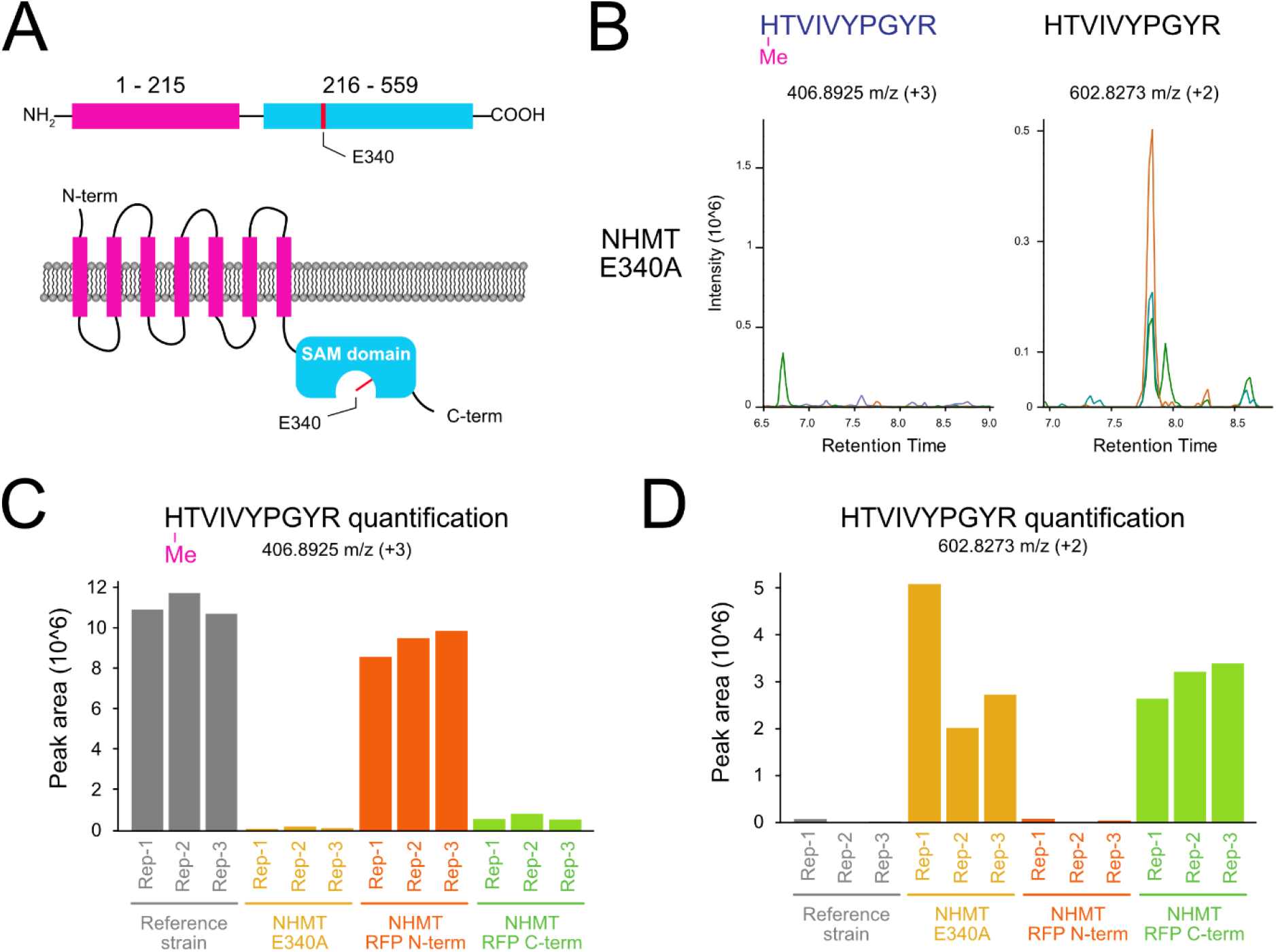
Functional and structural characterization of NHMT. **A)** Illustration of the NHMT candidate protein structure predicted by PSIPRED, showing the transmembrane domains (pink) and the soluble catalytic C-terminal (blue), where the targeted mutation E340A is shown in red. **B)** PRM chromatogram of the N-terminal histidine methylated and unmethylated HTVIVYPGYR peptide with an NHMT point mutation strain in the SAM catalytic domain at position E340A. **C)** Relative quantification of the N-terminal methylation histidine peptide ([meth]HTVIVPGYR) for the reference strain, the candidate with the E340A mutation on NHMT, and the NHMT candidate carrying an mRFP-tag at the N-terminal or the C-terminal. **D)** Relative quantification as in panel C of the unmethylated counterpart peptide HTVIVYPGYR.

To investigate the structural features and subcellular localization of NHMT, we expressed different NHMT constructs in *A. nidulans* with an mRFP fluorescence tag fused to the N-or C-terminal. PRM analysis of the different mRFP-expressing strains revealed that N-terminal histidine methylation was detectable for the N-terminally mRFP-tagged NHMT, but not with a C-terminal mRFP tag (Figure 3C, D, Supplementary Table S5). This suggests that the C-terminal mRFP tag interferes with the NHMTs catalytic activity.

### *In silico* analysis of NHMT agrees with the assignment of catalytic activity

Automated annotation of the gene product suggests that NHMT belongs to the spermidine synthase family of proteins (*27*). This is a subclass of the aminopropyl transferase family that catalyse the reaction between decarboxylated SAM and putrescine to form spermidine or longer polyamines (Supplementary Figure S3A) (*28, 29*). To establish NHMT as a putative methyltransferase and not spermidine synthase, the amino acid sequence of the soluble catalytic domain of NHMT was BLAST searched against the SwissProt database of manually curated sequences. This resulted in 48 protein sequences with predicted spermidine synthase and methyl transferase activities including histamine methyl transferases that similarly methylate the nitrogen atom of imidazole rings. Multiple alignments of these sequences displayed strong conservation at the SAM binding positions including the SAM binding glutamine 340 within NHMT that enabled us to generate a catalytic dead enzyme. However, the alignment also revealed a single amino acid variance within a conserved motif that separates spermidine synthases from other methyl transferases. Specifically, spermidine synthases contain either an aspartic acid (D) or glutamic acid (E) in the GxG(D/E)G motif located within the SAM binding and catalytic pocket (Figure 4A, Supplementary Figure S3D) that is required for spermidine synthase activity (*28, 30, 31*). However, isoleucine at residue position 322 (I322) replaces the D or E in this motif within NHMT (Figure 4A, Supplementary Figure S3). Other methyl transferases similarly lack a D or E in the motif, separating them from spermidine synthases in our analysis.

**Figure 4.**
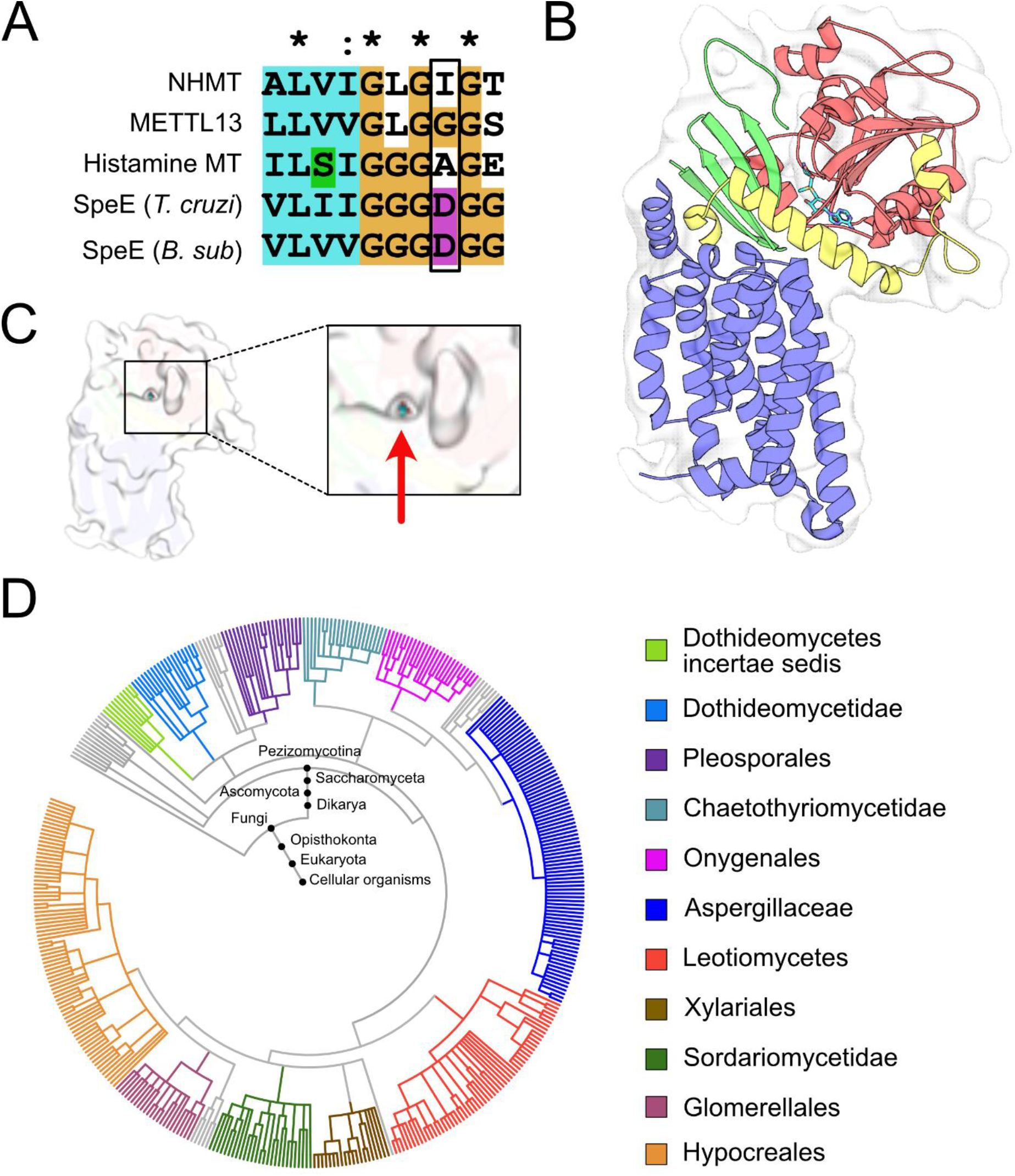
Motif and structure and activity prediction from the sequence of NHMT. **A)** Alignment of representative sequences, focused on the spermidine synthase motif. (Human eEEF1 n-terminal methyl transferase (METTL13), Human histamine n-methyl transferase, *Trypanosoma cruzi* spermidine synthase (SpeE *T. Cruzi*) and *Bacillus subtilis* spermidine synthase (SpeE *B. sub*). The residues corresponding to I322 are marked with a black box. The lack of an acidic group in this position shows that NHMT is not a spermidine synthase (also see Supplementary Figure S3C, D). **B)** The AlphaFold2 model of NHMT predicts four structural elements: 7TM (blue), four stranded anti-parallel beta-sheet (green), Rossmann-like domain (red) and a helical extension (yellow). **C)** The surface of the model is visualized with a white and slightly transparent rendering, and shows an opening in the structure that exposes the SAM molecule in the active site (marked by a red arrow). **D)** Phylogenetic analysis of the organisms with sequence similarity to NHMT 7TM (1-250). Major genera are highlighted in different colors. iTOL (*32*) tool was utilized to generate the phylogenetic tree.

### The transmembrane region of NHMT are predicted to be unique

The AlphaFold2 predicted structural model of NHMT includes a Rossmann-like domain (Figure 4B), a beta sheet rich region, and a unique C-terminal extension that wraps around the ectodomain that is ultimately buried in the transmembrane domains. The structural model predicts NHMT to contain an intermediate sized substrate-binding cavity required to contain the co-factor SAM and the substrate n-terminal histidine, which is in agreement with its NHMT activity (Figure 4C). The seven transmembrane helices (M1-M7) previously predicted with MEMSAT (Supplementary Figure S2) were also predicted by the AlphaFold2 model (Figure 4B). The topological orientation of the NHMT seven helices form a compact core, with hydrophobic residues facing outwards as one would expect for the membrane embedded domain with an approximate width of 40 Å, indicative of complete membrane spanning (Figure 4B). Searching the AlphaFold2 predicted 7TM domain model against the PDB25 DALI database (*33*) resulted in only weak hits with no 7TM proteins among them (Supplementary Figure S4C). Although G protein-coupled receptors are typically synonymous with 7TM domain, the helical organisation of the 7TM helices in NHMT show no structural or sequential similarity to the GPCRs.

Surprisingly, UniProt BLAST search of the 7TM domain (1-215) results in 452 matches to homologous 7TM domains containing proteins exclusively to organisms within the Ascomycota subphylum *pezizomycotina* (with the exception of one unclassified fungal organism), which contains the filamentous Ascomycetes including *Aspergillus* (Figure 4C, Supplementary Figure S4B). The analysis revealed ~97% of the matches are single copy genes within their respective organisms. Moreover, all 452 transmembrane domain sequences were linked to soluble domains that contain a highly conserved segment 389-394 uniquely found in proteins with the NHMT 7TM domain (Supplementary Figure S4B). This analysis thus revealed that the NHMT represents a hitherto uncharacterized protein family containing a 7TM domain linked to a methyl transferase domain that likely catalyzes methylation of n-terminal histidine in filamentous fungi.

### NHMT resides in the endoplasmic reticulum and methylates N-terminal histidines of secreted proteins

The LPMOs identified to date are typically secreted outside of the cell by their host organism. Furthermore, all four proteins detected with N-terminal histidine methylation in our proteomics dataset are encoded with a signal peptide, indicating processing through the endoplasmic reticulum (ER). We therefore hypothesized that NHMT must be located in the ER to methylate its substrates. To test this, we generated a truncated variant of NHMT lacking the transmembrane region (positions 1-224); the expression of this variant abolished the N-terminal histidine methylation capacity of the enzyme, indicating that the transmembrane region is critical for its specific activity (Figure 5A, B, Supplementary Table S5). We also performed co-localization cellular imaging by fluorescence microscopy with the addition of an mRFP tag to the C-or N-terminal of full length NHMT alongside a positive ER marker, the fluorescent organelle probe DiOC₆(*34*) (Figure 5C). We utilized cytosolic mRFP as a negative ER membrane control (Figure 5D), and a mannosyltransferase protein known to localize to the ER (AN10118, UniProt entry C8VRA6) involved in protein glycosylation was tagged with mRFP and used as a positive ER localization control(*35*). The imaging analysis revealed higher co-localization of all mRFP-tagged NHMT constructs with the ER membrane control protein, confirming subcellular localization in the ER (Figure 5C, D).

**Figure 5.**
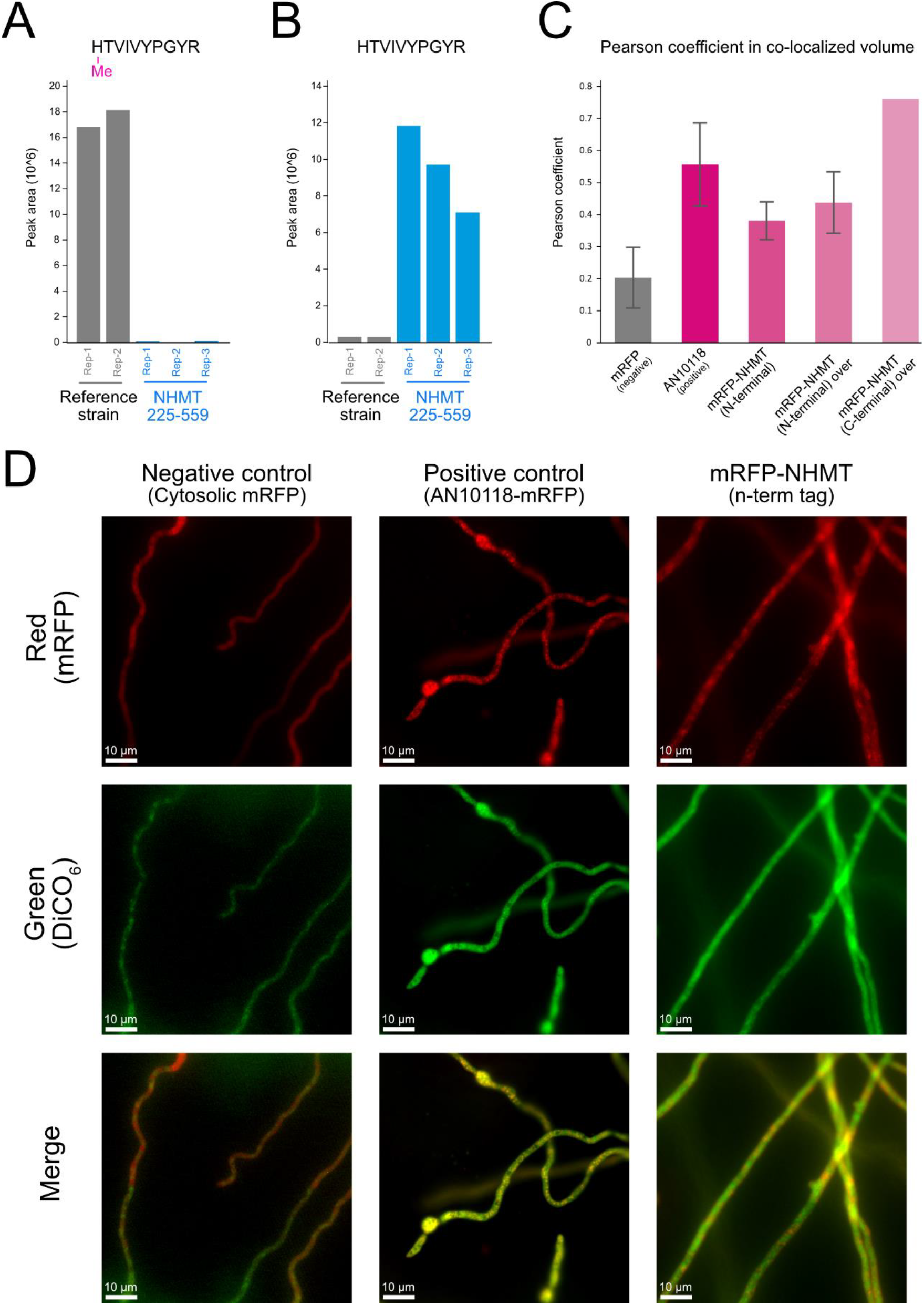
Impact of NHMT truncation on activity and NHMT ER localization. **A)** PRM quantification of methylated HTVIVPGYR due to truncation of the transmembrane domain of NHMT. **B)** PRM quantification of unmethylated HTVIVPGYR after NHMT truncation. **C)** Microscopy quantification of NHMT ER co-localization of different strains as indicated by the Pearson correlation. Error bars indicate the standard error of the mean. **D)** Co-localized image of negative control cytosolic mRFP, positive control endogenous mRFP tagged ER protein C8VRA6, mRFP-NHMT (N-terminal tagged) expressing cells. Two different channels: red (mRFP), green (DiCO_6_), and merged are shown, yellow indicates co-localization in the merged channel.

### Recombinant methylation of LPMOs by NHMT in the yeast *Komagataella phaffii*

To test NHMT activity and determine its biotechnological potential for recombinant N-terminal histidine methylation, we co-expressed NHMT together with a fungal AA9A LPMO from *Lentinus similis* (LsAA9A, UniProt accession A0A0S2GKZ1) or a bacterial AA10 LPMO from *Thermobifida fusca* (TfAA10A, UniProt accession Q47QG3) in the yeast protein production host *Komagataella phaffii* (formerly *Pichia pastoris*). The full-length *nhmT* sequence from *A. nidulans* was codon-optimized and expressed without introns in *K. phaffii*. We tested three different secretion peptide sequences for protein translocation through the ER using LsAA9A as the methylation substrate (Figure 6A). These were the native LsAA9A signal peptide, the α-amylase (Amy^SP^) secretion sequence from *A. niger* termed “Amy” (*36*), and the alpha-mating factor (α-MF) secretion signal leader peptide from *S. cerevisiae* (*37, 38*). LsAA9A expression and activity were confirmed both with and without co-expression of NHMT (Figure 6B). LsAA9A was secreted into the supernatant with all three signal peptides (Supplementary Figure S5, S6, S7), however correct signal peptide processing was only observed and confirmed with LC/MS/MS when using the native and the Amy signal peptide (Figure 6C, Supplementary Table 6, Supplementary Figure S8). Additionally, we did not detect histidine methylation by MS when the peptide was not cleaved at the correct position with the α-MF as the signal peptide sequence, suggesting that N-terminal histidine methylation occurs only after precise signal peptide cleavage. We quantified the degree of N-terminal histidine methylation to be ~30% of the identified tryptic LsAA9A N-terminal peptide when co-expressed with NHMT (Figure 6C, D). TfAA10A was similarly found to be secreted into the supernatant when the sequence was expressed with the Amy signal peptide (Supplementary Figure S9). Furthermore, we observed an average of 15% N-terminal histidine methylation of TfAA10A (Figure 6E, Supplementary Table S7), demonstrating that it is possible to engineer a heterologous expression host for specific methylation of N-terminal histidines on different protein targets.

**Figure 6.**
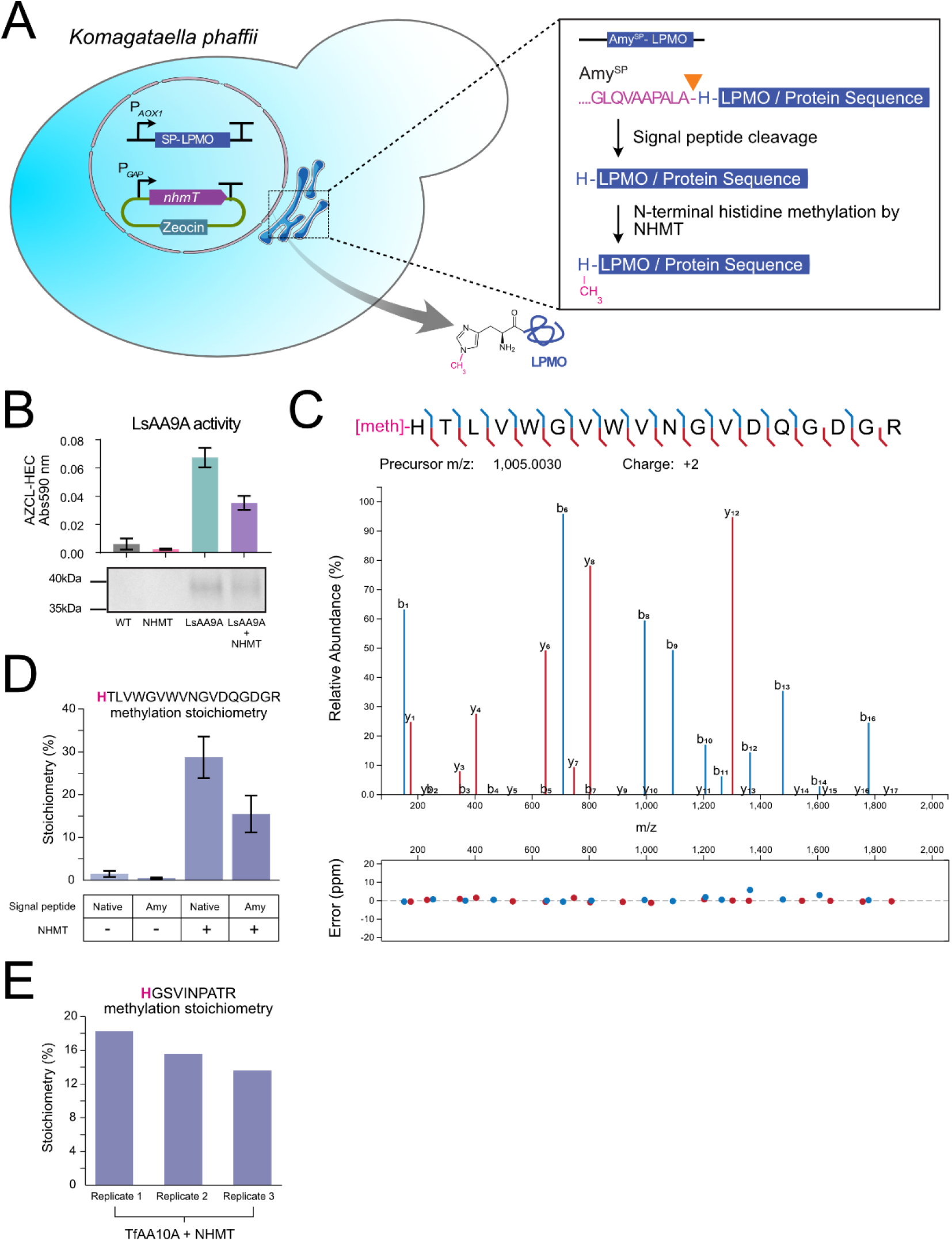
Co-expression strategy of *nhmT* and recombinant N-terminal histidine methylation. **A)** Schematic of heterologous *nhmT* co-expression in *K. phaffii* for generating recombinant proteins. A DNA construct encoding an LPMO is inserted into the genome and expressed under the strong methanol-inducible promoter p*AOX* together with *nhmT* expressed from an episomal plasmid. The signal peptide ensures translocation into the ER prior to secretion into the extracellular medium. **B)** LsAA9A activity (upper graph) as measured by AZCL-HEC assay (see methods). All samples were analysed in biological triplicates; error bars indicate standard deviations. SDS-PAGE (lower figure) of the LsAA9A secreted into the supernatant of one of the biological replicates. **C)** MS/MS spectra annotation(*39*) of a high-confidence identification of N-terminal histidine methylated tryptic peptide of LsAA9A with full sequence coverage and low fragment mass error. **D)** N-terminal methylation stoichiometry calculation via precursor quantification of the methylated and unmethylated N-terminal tryptic LsAA9A peptide HTLVWGVWVNGVDQGDGR. LsAA9A is secreted into the medium in the presence or absence of the NHMT. All samples were analysed in biological triplicates; error bars indicate standard deviations. **E)** N-terminal methylation stoichiometry calculation via precursor quantification of the methylated and unmethylated N-terminal tryptic TfAA10A peptide HGSVINPATR upon co-expression of TfAA10A and NHMT in *K. phaffi*.

## DISCUSSION

In this study, we have identified the elusive gene *nhmT* (AN4663) encoding the NHMT enzyme solely responsible for N-terminal histidine methylation of LPMOs. This was accomplished through a combination of in-depth proteomics and CRISPR/Cas9 facilitated gene knockouts. Consequently, we identified close to 7000 *A. nidulans* protein groups from the proteomics analysis, generating the largest dataset of filamentous fungal proteins to date. Quantitative proteomics analysis further enabled us to narrow down the list of potential NHMT candidates that were selected for CRISPR/Cas9 gene knockouts. We developed a targeted proteomics assay to determine the effect of these knockouts and ultimately identified the putative NHMT candidate.

We determined that *nhmT* encodes an NHMT enzyme that shares the seven-β-strand fold with the human histidine methyltransferases METTL9 and METTL18 as well as the human N-terminal methyltransferase METTL13. NHMT also contains an integral membrane region with a potentially novel seven-transmembrane with little structural similarities when compared to known membrane proteins the PDB database. The transmembrane is required for NHMTs specific activity. The 7TM region is unique to filamentous fungi and genes with similar sequence similarity are found primarily as single copies in the genomes of filamentous fungi, suggesting distinct phylogeny of the enzyme.

Loss of the N-terminal histidine methylation due to transmembrane truncation could be due to the loss of co-localization with its substrate proteins that require signal peptide cleavage through the ER for secretion. Co-localization imaging suggested that NHMT is indeed located in the ER, providing further support of this theory. However, it is also plausible that the transmembrane region of NHMT is important for trafficking to the ER membrane, correct protein folding and putative oligomeric assembly (*40*). The transmembrane region might harbour sites for lipid specific interactions, which be necessary for its specificity towards N-terminal histidine residues. Furthermore, we observed a significant reduction (but not loss) of methylation activity upon the addition of an mRFP tag at the C-terminal of the NHMT, suggesting loss of catalytic function or substrate recognition, via either structure inhibition or substrate blockage.

The discovery of NHMT enabled co-expression with LPMOs in *K. phaffii*, leading to the demonstration of recombinant N-terminal histidine protein methylation in a heterologous host that is not a filamentous fungus. Future efforts will be directed towards optimizing the co-expression of the NHMTs and the targeted substrates to increase methylation stoichiometry. The findings presented here demonstrate a potential to generate recombinant proteins with methylated N-terminal histidines, which could enable further investigations into the role of this modification in LPMO function and activity. N-terminal histidine methylation could also have potential applications beyond LPMOs, in the production of different classes of pharmaceutical proteins and peptide hormones such as incretins; where methylation of the terminal histidine of GLP-1 for example, may incur protease protection to plasma dipeptidyl peptidase-4 (*41, 42*).

This study also highlights the utility of combining large-scale quantitative proteomics with a systematic CRISPR editing screen of specific enzyme classes for discovering novel biology in organisms such as the filamentous fungus *A. nidulans*. Filamentous fungi play a crucial role in the degradation of biopolymers and thereby the recycling of organic matter, enabled by the secretion of a battery of enzymes. These enzymes are best known for their roles in biomass degradation that has been exploited in biotechnological and industrial settings (*10*). Exploring and characterizing fungal enzymes will thus affect a broad number of bio-based industries. The results presented here pave the way for further proteomics investigations into the diverse phyla of the fungal eukaryotic kingdom, aimed at the discovery of novel PTMs that may lead to the discovery of unique protein attributes.

## ABBREVIATIONS

LPMO: lytic polysaccharide monooxygenase.
NHMT: N-terminal histidine methyltransferase.
PRM: parallel reaction monitoring.
LC: liquid chromatography.
MS: mass spectrometry.
LFQ: label-free quantification.
SAM: S-adenosylmethionine.
ER: endoplasmic reticulum.
m/z: mass-to-charge ratio.
PTM: post-translational modification.

## ACKNOWLEDGEMENT

We would like to acknowledge Antonios Georgantzoglou and Jutta Maria Bulkescher for their assistance in fluorescence microscopy and image analysis. We thank Charles C. Lee for providing us with the episomal plasmid pBGP1 for episomal expression in *K. phaffii*. We thank Helle Juel Martens assistance with flourescene microscopy of *A. nidulans*. This work was supported by the Novo Nordisk Foundation (NNF14CC0001 and NNF17SA0027704),

## AUTHOR CONTRIBUTIONS

K.S.J. proposed the study. T.S.B., J.B.H., and J.V.O. conceived the scientific strategy. T.S.B. designed the proteomic strategy and analysed all proteomics-related samples and corresponding data. J.B.H. initiated the development of *A. nidulans* CRISPR/Cas9 experiments. T.S.B. and J.B.H. designed the knockout candidate list. J.B.H. and J.L.S. designed the *A. nidulans* engineering strains and expressed the strains for the proteomics and imaging analysis. C.H.R. and M.H.H.N. contributed to the strain engineering of the knockout strains in *A. nidulans*, designed and engineered *K. phaffii* for the methylation of LPMOs. S.B. performed the protein sequence alignments and structural analysis. J.P.M. performed AlphaFold2 predictions and protein modelling. T.S.B. and J.V.O. conceptualized and established the manuscript. All authors contributed to the final version of the manuscript.

## MATERIALS AND METHODS

All strains, vectors, fragments, primers, and DNA sequences used in this study are listed in supplementary information (Table S3, S4, S5, S6, and Note S1-S3).

### Strains and cultivation media

The strain NID2531 (*argB2, veA1, nkuA*Δ, Supplementary Information, Table S3) was used as the reference strain and subsequent studies and host for all gene deletions and insertions. The strain contained a mutation in the gene encoding ornithine transcarbamylase (*argB*2) of the arginine biosynthesis pathway, which is utilized as selection in transformations via the auxotrophic growth requirement of arginine. Deletion of *nkuA*Δ eliminates to a large extent the activity of the non-homologous end-joining DNA repair mechanism and forces repair to take place through homologous recombination to promote gene targeting(*22*). NID2531 is derived from the wild-type strain FGSC A4, which was used for the initiating *Aspergillus nidulans* proteome experiment. Spore production for inoculations and strain validation were produced in solid minimal medium (MM: 2% agar, 1% glucose, 1 x nitrate salt solution, 0.001% thiamine, 1 x trace metal solution) (*43, 44*), which was supplemented with 4 mM L-arginine when required.

The parent strain used for engineering and protein production was *Komagataella phaffii* GS115 (Thermo Fisher Scientific, Waltham, MA, USA), which has a mutation in the *his4* gene, rendering the strain unable to produce histidine. All plasmids employed for genomic integration contain the HIS4 gene for complementation of the *his4* gene in the parent strain. Transformants are plated on selective medium lacking histidine.

### DNA fragment and plasmid constructions in *A. nidulans* and *K. phaffii*

Fragments constructed by PCR were made as described previously(*22*) and CRISPR-Cas9 vectors were assembled by USER fusion(*45*). The vectors for *K. phaffii* expression were constructed by LyGo cloning as described in Hernández-Rollán *et al*. (*36*), and by USER fusion as described previously(*46*). Primers and synthetic gene fragments were purchased from Integrated DNA Technologies (IDT, Coralville, IA, USA), and are listed in Tables S6. Synthetic gene AN4663 used for expression in *K. phaffii* is listed in Supplementary Notes S3. All PCR fragments for cloning and plasmids were purified using NucleoSpin Gel and PCR Clean-up kit (Macherey-Nagel) and GenElute™ Plasmid Miniprep Kit (Merck), respectively, and the vector assemblies were confirmed by sequencing (Eurofins Genomics, Germany).

All deletions and codon substitutions in *A. nidulans* were enabled by the oligonucleotide-mediated gene-editing procedure described previously (*22*), except for the length of the gene-editing oligonucleotides which was 60 nucleotides instead of 90. All CRISPR/Cas9-vectors and expression vectors were assembled via uracil excision-based cloning (USER), as described previously(*22*). All CRISPR/Cas9 vectors were based on and pFC331 as the recipient vector(*21*), and were composed by one or more single guide RNAs flanked by tRNAs. All PCRs for DNA-fragment construction in *A. nidulans* engineering were done as described previously (*22*). The expression vector for insertion into integration site 5 (IS5) was established in pU0002 as described previously(*47*) with a recreated PacI/Nt.BbvCI USER cassette between the targeting sequences of 1.0 kb matching IS5 for cloning of promoter Ptef1, gene fragment, and terminator Ttef1.

For heterologous expression in *Komagataella phaffii*, genes for LsAA9A and TfAA10A were cloned according to the LyGo cloning protocol and integrated into the genome of *K. phaffii*. The native DNA sequence of LsAA9A followed by a TEV cleavage recognition site and a His purification tag was cloned into three different constructs carrying three different signal peptides for secretion in the yeast *K. phaffii*. The following signal peptides were selected: α-MF signal peptide resulting in the vector named pLyGo-*Kp*-1-LsAA9A, Amy^SP^ signal peptide from *A. niger* resulting in the vector pLyGo-*Kp*-2-LsAA9A, and the native *L. similis* signal peptide of LsAA9A resulting in the vector called pPIC9K-Native^SP^-LsAA9A. The LsAA9A native signal peptide and cloned into the vector pLyGo-*Kp*-2-LsAA9A using USER cloning in which both the forward (5990) and the reverse primer (5991) contained the native signal peptide, resulting in the vector pPIC9K-Native^SP^-LsAA9A. The native DNA sequences without their native signal peptides of TfAA10A from *Thermobifida fusca* (UniProt: Q47QG3) was ordered as synthetic gene fragments from Integrated DNA Technologies (IDT, Coralville, IA, USA) with SapI restriction enzyme sites compatible with LyGo cloning. The gene fragment was cloned by LyGo cloning into the vector pLyGo-*Kp*-2 with Amy^SP^, resulting in the vector pLyGo-*Kp*-2-TfAA10A. The signal peptide Amy^SP^ was chosen to direct the proteins to the ER as it was predicted to be cleaved at position His1 for both proteins according to SignalP (*48*). All vectors were cloned and propagated in *E. coli* DH5a, mini-prepped, and their sequences confirmed by Eurofins Genomics (Ebersberg, Germany).

The synthetic DNA sequence of AN4663 was purchased from Integrated DNA Technologies (Coralville, IA, USA) without introns codon-optimized and cloned into the episomal vector pBGP1 (a kind gift from Charles Lee) (*49*) using uracil excision-based cloning as described previously (*46*) using the primer pair 5701 and 5702. The expression of the AN4663 in the episomal plasmid pBGP1 is driven by the strong constitutive promoter pGAP. The resulting plasmid was verified by DNA sequencing.

### Competent cells, transformation, and strain validation

Protoplastation for *A. nidulans* was performed as previously (*50*). The transformations were made in gently thawed protoplasts(*22*) after which the strains were incubated at 37°C and validated by diagnostic PCR as described previously(*21*). Each transformation protoplasts were mixed with 1.5 μg of CRISPR vector and used in combination with either 2 μg of a linear double strand DNA for homologous recombination or 20 μL of 100 μM stock solutions of oligo nucleotides in a total volume of 150 μL of PCT buffer (50 % w/v PEG8000, 50 mM CaCl2, 20 mM Tris, 0.6 M KCl, pH 7.5). The mix was incubated for 10 minutes at room temperature, followed by addition of 250 µl of transformation buffer (1.2 M sorbitol, 50 mM CaCl2· 2 H2O, 20 mM Tris, 0.6 M KCl, pH 7.2), and plated on transformation media (1M sucrose, 2% agar, 1 x nitrate salt solution, 0.001% thiamine, 1 x trace metal solution plates). All transformation plates were incubated at 37°C. Resulting transformants were examined via diagnostic tissue-PCR.

For *K. phaffii*, all vectors were linearized before genomic integration using the primers 4917 and 4918 and the linearized fragments were confirmed by electrophoresis prior to electroporation. PCR fragments were further subjected to PCR clean-up using PCR clean-up kit (Macherey-Nagel™). Electrocompetent cells were produced according to the Invitrogen Pichia Expression Kit (Catalogue no. K1710-01). In detail, a starting culture of the desired strain was grown in 5 mL yeast extract peptone dextrose (YPD) medium at 30°C overnight. Next day, 0.1 mL was used to inoculate a fresh culture of YPD and incubated at 30°C overnight until an optical density measured at a 600 nm wavelength (OD_600_) of 1.3 was obtained. On the following day, the culture was centrifuged at 1500 g at 4°C and the cells resuspended in 500 mL ice-cold water. The process was repeated with 250 mL ice-cold water, and once more with 20 mL of an ice-cold sorbitol solution (1 M). After final centrifugation, the cells were resuspended in 1 M sorbitol solution to a final volume of 1.5 mL. Approximately 1 µg of PCR product was used in combination with 80 µL of freshly produced electrocompetent cells (GS115) for electroporation. The cells and the DNA were incubated on ice for 5 minutes prior to electroporation. For electroporation, a 0.2 cm gap sterile electroporation cuvette was used, and the cells were electroporated using the following parameters: (voltage (V) 1500, capacitance (µF) 25, and resistance (Ω) 200). Right after electroporation, 1 mL of 1 M ice-cold sorbitol was added to the cuvette, and the entire cell pulsed was then plated on minimal dextrose medium (MD), and incubated upside down at 30° for 3-4 days until colonies had formed. The identity of the transformants was confirmed by colony PCR. Colonies grown on MD plates were picked and resuspended in 50 µL of 20 mM NaOH and boiled at 99°C for 25 min. The mixture was centrifuged for 1 minute at 11,000 g to remove cellular debris, and 5 µL of the supernatant was used as a template for the PCR reaction using OneTaq® 2X Master Mix with Standard Buffer (New England Biolabs) primers 4925 and 4926.

Correct clones producing LsAA9A and TfAA10A were selected for small-scale (in 24-well plates) screening of protein production, and the best producers were then used to make a second batch of electrocompetent cells for the introduction of the episomal plasmid containing AN4663. Selected strains of LsAA9A and TfAA10A expression were made electrocompetent using the method described above, and 1 µg of the episomal plasmid pBGP1-AN4663 was electroporated as described above. After electroporation, 1 M ice-cold sorbitol solution was added to the cuvette, and the mixture was recovered at 30°C for 2 h with vigorous shaking. The entire culture was then plated on selective YPD plates containing 100 µg/mL Zeocin™ and incubated for up to a week at 30°C until colonies appeared.

### Co-expression of LsAA9A and TfAA10A together with AN4663

Single colonies of the desired strain were used to inoculate 10 mL of Buffered Glycerol Complex Medium (BMGY) (Pichia Expression Kit, Life Technologies, Carlsbad, CA, USA) and grown at 28°C for two days with shaking at 250 rpm. From the saturated culture, a starting culture with an OD600 of 1.0 was prepared in 25 mL buffered methanol complex (BMMY) medium, sometimes supplemented with Zeocin (100 µg/mL) for the strains carrying the pBGP1 plasmid. Baffled glass flasks were used, and the cultures were incubated at 28°C with shaking at 250 rpm. Expression was maintained by the addition of 1% methanol every day for an additional four days. OD_600_ measurements were carried out daily and the cultures were plated on YPD-agar with 100µg/ml Zeocin at the end of expression to check for the presence of the pBGP1 plasmid containing AN4663. On the final day, the cells were collected by centrifugation at 5000 g for 20 minutes, and the supernatant was subjected to filter sterilization. To check for the secretion of LsAA9A and TfAA10A, 5 µL of the supernatant was loaded into a 4−20 % Mini-PROTEAN-TGX gel (BioRad, Hercules, CA, USA), run at 165 V for 50 minutes. After the run, gels were stained with InstantBlue Protein Stain (Expedeon Inc., Inc., San Diego, CA, USA) for one hour and destained over night with demineralized water.

### AZCL-HEC assay

The activity of secreted LsAA9A from *K. phaffii* was determined by mixing 1 mg/mL AZCL-HEC substrate (Megazyme, County Wicklow, Bray, Ireland), with 1 mM ascorbic acid (Sigma-Aldrich, Saint Louis, MO, USA), 100 μM copper sulphate (Sigma-Aldrich), and the volume was adjusted with 100 mM sodium acetate (pH 5) (Sigma-Aldrich). 100 µL of the secreted LsAA9A sample was mixed with 400 μL of the AZCL-HEC substrate reaction and incubated at 50°C with shaking at 1500 rpm for 1 hour. The samples were centrifuged to remove the AZCL-HEC substrate, and the absorbance was measured at 590 nm.

### Fluorescence microscopy and image analysis

Fresh *A. nidulans* spores suspensions (10 µL of ~10^5^ spores per mL) were inoculated on glass slides with 0.5 mL solid MM (as described above) with 4 mM L-arginine and incubated for 20 hours in petri dishes in micro-perforated bags at 37 °C. A cover slide and immersion oil were applied and the ER was labelled with 10µM 3,3′-Dihexyloxacarbocyanine iodide (DiOC_6_) for 20 minutes. Images of *A. nidulans* were acquired with a Leica DMI6000 widefield microscope, equipped with an 63×1.40 OIL HC PL APO objective and operated with LAS X (version 3.3.3) software. Three images of mRFP tagged proteins, green mitochondrial membrane dye (DiOC_6_) and bright field were acquired for each sample. Collected images were processed using the Imaris software version 9.8.2 (Bitplane AG, Zürich / Oxford Instruments, Abingdon, Oxfordshire, England). Background subtraction was performed followed by manual segmentation of *A. nidulans* filaments and saved as surfaces for each image and co-localization of mRFP and DiOC_6_ labelled components were determined using Imaris Coloc tool. Intensity threshold of 1000 and 600 were used respectively for the mRFP (red) and DiOC_6_ (green) channels in all images.

### Proteomics sample lysis and preparation

All fermentations for proteomics analysis were inoculated with a fresh spore suspension in 20 mL liquid media to a concentration of 10^6^ spores per mL in falcon mini bioreactors. Three different liquid media for *A. nidulans* were utilized for the fermentations, these include potato dextrose media containing 39 g/L potato dextrose, minimal media with 1% glucose, or cellulose minimal media (0.5% cellulose, 1% glucose). All media contained 1x nitrate salt solution, 0.001% thiamine, and 1 x trace metal solution supplemented with 4 mM L-arginine. Fermentations were carried out for 72 hours at 37°C at 45° degree angle with 180 RPM shaking. Biomass was filtered and biomass were washed with Phosphate-buffered saline and frozen at −80°C. Five mL of *A. nidulans* cell biomass was pelleted by centrifugation and washed with cold PBS. Lysis buffer (2% Triton X-100, 1% SDS, 100 mM NaCl and 10 mM EDTA) was added to each sample and vortexed, followed by incubation at 95°C for 10 minutes and then vortexed again. 1.5 mL of the resulting solution was transferred to 2 mL bead beating tubes containing 1.4 millimetre Zirconium oxide beads. The tubes were subjected to beating using a Precellys 24 homogenizer (Bertin Technologies, Montigny-le-Bretonneux, France) at 6800 rounds per minute for 30 s, followed by sample cooling for 60 s; this was repeated 4 times for a total of 5 times. The resulting liquid was transferred to new 1.5 mL tubes and sonicated using a 2 millimetre sonication tip at 100% power, with 3 s ON and 1 s OFF, for a total sonication time of two minutes. The resulting liquid was centrifuged at 20,000 g for 10 minutes at 4°C, and the resulting supernatant transferred to new 1.5 mL tubes. The protein concentration in the samples was determined using a tryptophan assay(*51*).

Cysteine residues of the resulting protein lysate were reduced with 5 mM tris(2-carboxyethyl)phosphine and alkylated with 5 mM chloroacetamide for 30 minutes at room temperature. Protein lysate was prepared for protease digestion with protein aggregation capture (PAC) as described previously(*52*) using magnetic hydroxyl beads. Once the proteins had aggregated on the beads, 50 mM HEPES buffer (pH 8.5) was added. This was followed by the addition of Lys-C protease at a ratio of 1:200 (to total lysate protein) and digestion at 37°C for 4 hours, followed by the addition of trypsin at a ratio of 1:50, and digestion at 37°C overnight. Samples were acidified to quench the protease reaction by the addition of trifluoroacetic acid (TFA) to a final concentration of 2%.

The resulting peptide mixture was prepared for downstream LC/MS/MS analysis with hydrophobic solid-phase extraction using C18 Waters Sep-Paks (Milford Massachusetts, USA). Briefly, Sep-Paks were prepared by the sequential addition of acetonitrile and 0.1% TFA solution, followed by the addition of the acidified peptide mixture. The samples were then washed with 0.1% TFA and eluted into new tubes using 50% acetonitrile solution with 0.1% TFA. Acetonitrile was evaporated from the peptide solution using an Eppendorf Concentrator Plus speedvac operated at 60°C. Dried peptides were reconstituted in 0.1% TFA solution, and the peptide concentration was determined using a Thermo Fisher NanoDrop spectrophotometer.

For samples analysed by in-gel proteomics analysis, sample preparation was performed as previously described(*53*).

### TMT labelling

Peptides were prepared for labelling using the PAC protocol described above. Briefly, 20 µg of protein was protease-digested on-bead overnight in 50 mM HEPES buffer (pH 8.5). The resulting peptide mixtures were transferred to new tubes and acetonitrile was added to a final concentration of 50%. Tandem mass tag 11-plex reagents (Thermo Fisher Scientific, Waltham, Massachusetts, USA) were added to individual sample and incubated for 60 minutes at room temperature. The resulting reaction was quenched by the addition of 5% hydroxylamine for 15 minutes, followed by the addition of TFA to a final concentration of 1%. The labelled samples were pooled and mixed, followed by the evaporation of acetonitrile with a speedvac (see above). The resulting peptide pellet was reconstituted in 50 mM ammonium bicarbonate buffer (pH 8.5) and prepared for offline fractionation.

### High pH reversed-phase offline peptide fractionation

A mixture of 200 µg of the peptides reconstituted in 50 mM ammonium bicarbonate buffer (pH 8.5) was injected, fractionated and collected using a Thermo Scientific UltiMate 3000 high-performance liquid chromatography (HPLC) system. The system was operated at 30 µL/min, and peptides were separated on a Waters Acquity CSH C18 1.7 μm 1 × 150 mm C18 column. Aqueous buffer A (5 mM ammonium bicarbonate) and buffer B (100% acetonitrile) were used for the gradient, which consisted of increasing B from 8% to 28% in 62 minutes, followed by an increase to 60% B, and another ramp to 70%, where it was maintained for 8 minutes, followed by ramp back down to 8% B, where it was maintained for 10 minutes. For TMT-labelled peptides, the gradient was initially increased to 30% B, followed by a ramp to 65% B and rapid increase to 80% B, where it was maintained for 7 minutes. The total fractionation time for both methods was 87 minutes, and a total of 46 fractions were collected. Formic acid (10%) was added to each fraction, and the samples were dried to completeness using the speedvac. Dried fractions were reconstituted in 40 µL 0.1% formic acid and 20 µL of each fraction was loaded onto Evotips (EvoSep, Odense, Denmark) according to the manufacturer’s instructions, and prepared for MS analysis.

### Liquid chromatography coupled to mass spectrometry

Single-shot label-free quantification (LFQ) analysis of protein digests from *A. nidulans* cells grown on different carbon sources were analysed using an EASY-nLC 1200 system (Thermo Fisher Scientific) coupled to the MS with 0.1% formic acid as buffer A and 80% acetonitrile with 0.1% FA as buffer B. 500 ng of peptide mixture was injected onto a homepacked C18 column (15 cm x 75 μm inner diameter) packed with 1.9 μm C18 beads (Dr. Maisch GmbH, Entringen, Germany). The peptide mixture was separated using a gradient of 0 to 25% B in 60 minutes, after which it was increased to 40% B in 15 minutes. The gradient was further increased to 80% B in 5 minutes, where it was held for an additional 5 minutes, followed by a ramp down to 5% B in 3 minutes where it was held for 2 minutes for equilibration. The total runtime was 90 minutes. A column temperature of 50°C was applied during all the runs using a column oven (PRSO-V1, Sonation, Biberach, Germany).

An EvoSep One nano-LC system (EvoSep, Odense, Denmark) was coupled to MS for peptide sequencing analysis of individual fractionated samples (TMT labelled and unlabelled) and PRM analysis. Two methods were utilized for all samples: either the 60 samples per day, resulting in a total runtime of 21.5 minutes, or 30 samples per day, with a total runtime of 44 minutes. Buffer A, consisting of 5% acetonitrile with 0.1% formic acid, and buffer B, consisting of 0.1% formic in acetonitrile, were used for separation. A 15 cm x 150 μm inner diameter homepacked column with 1.9 μm C18 beads was heated to 60°C, as described above.

### Mass spectrometric analysis

The samples were analysed on either a Orbitrap Q-Exactive HF-X or Orbitrap Exploris 480 mass spectrometer (Thermo Fisher Scientific) operated in positive mode with 2 kV spray voltage and a capillary temperature of 275°C. An MS1 resolution of 120,000 and an MS2 resolution of 30,000 (with an automated gain control (AGC) fill time of 54 seconds) were used for data-dependent acquisition (DDA). A normalized collisional energy (NCE) of 28 and a quadrupole isolation window of 1.3 m/z were used for LFQ samples, while corresponding values of 35 NCE and 0.8 m/z isolation window were used for TMT-labelled samples.

Targeted analysis was performed using parallel reaction monitoring (PRM) with the Orbitrap Exploris 480 mass spectrometer. An MS1 scan with a resolution of 120,000 was performed prior to an unscheduled product ion scan that cycles through an inclusion list of peptide mass-to-charge (m/z) ratios provided for targeted analysis. Typically, two charge states for each peptide were targeted for PRM analysis in order to increase the specificity. An isolation width of 1.3 m/z, an MS2 resolution of 60,000, a NCE of 30, and maximum injection time of 100 ms were used for each target.

### Mass spectrometry data analysis

All DDA data were searched using MaxQuant software(*54*). *A. nidulans* samples were searched against the UniProt *Emericella nidulans* (taxonomy ID 227321, strain FGSC A4 / ATCC 38163 / CBS 112.46 / NRRL 194 / M139) reference proteome. Signal peptides predicted by SignalP(*48*) were removed from the leading sequence of all signal peptide containing proteins in the FASTA file prior to searching. Files were searched with “Trypsin/P” as the protease specificity, with a maximum of two missed cleavages, fixed carbamidomethyl modification on the cysteines, and variable modifications of oxidation (methionine), acetylation (protein N-terminal), methylation (histidine), phosphorylation (serine, threonine, and tyrosine), and deamidation (asparagine, and glutamine) were utilized. A methylated histidine-specific diagnostic ion of 124.0869 m/z was utilized to assist in the high-confidence localization of methylated histidines(*55*). The search mass tolerance was first set to 20 ppm, followed by 4.5 ppm after main search recalibration. The mass tolerance for fragment ions was set to 20 ppm. An Andromeda score cut-off of 40 was utilized with a 1% false discovery rate at all levels (peptide spectral matches, peptides, and proteins).

PRM and stoichiometry analysis were performed with the Skyline software(*56*). A spectral library was constructed in Skyline from MaxQuant-processed results from label-free samples to assist peak identification. The PRM results were quantified by summing the fragment peak areas (Supplementary table S5). Summed MS1 peak areas of the three isotopic parent peptide masses were utilized for quantitative comparison in the stoichiometry analysis.

### Downstream bioinformatics analysis

MaxQuant-processed results were analysed using the Perseus software(*57*). LFQ intensities reported in the proteingroups.txt output from MaxQuant were used for the quantitative analysis of label-free samples analysed with LC/MS/MS. Contaminant and reverse hits were removed from the list and the values were log2 transformed. Protein groups were removed if not observed in at least two replicates in one condition. Missing values were imputed using a width of 0.3 and downshift of 1.8. To determine regulated proteins, a two-sided t-test was performed using a permutation-based false discovery rate cut-off of 0.05 as truncation for significant hits. For TMT-labelled samples, reporter intensities were utilized instead of LFQ. Contaminants and reverse hits were similarly removed, and quantile normalization was performed on the reporter intensities followed by log2 transformation. A two-sided t-test was performed as described above to determine significant differences between proteins for LFQ and TMT experiments.

## REFERENCES

1. T. Jenuwein, C. D. Allis, Translating the Histone Code. Science. 293, 1074–1080 (2001).

2. A. W. Wilkinson, J. Diep, S. Dai, S. Liu, Y. S. Ooi, D. Song, T.-M. Li, J. R. Horton, X. Zhang, C. Liu, D. V. Trivedi, K. M. Ruppel, J. G. Vilches-Moure, K. M. Casey, J. Mak, T. Cowan, J. E. Elias, C. M. Nagamine, J. A. Spudich, X. Cheng, J. E. Carette, O. Gozani, SETD3 is an actin histidine methyltransferase that prevents primary dystocia. Nature. 565, 372–376 (2019).

3. E. Davydova, T. Shimazu, M. K. Schuhmacher, M. E. Jakobsson, H. L. D. M. Willemen, T. Liu, A. Moen, A. Y. Y. Ho, J. Małecki, L. Schroer, R. Pinto, T. Suzuki, I. A. Grønsberg, Y. Sohtome, M. Akakabe, S. Weirich, M. Kikuchi, J. V. Olsen, N. Dohmae, T. Umehara, M. Sodeoka, V. Siino, M. A. McDonough, N. Eijkelkamp, C. J. Schofield, A. Jeltsch, Y. Shinkai, P. Ø. Falnes, The methyltransferase METTL9 mediates pervasive 1-methylhistidine modification in mammalian proteomes. Nature Communications. 12, 891 (2021).

4. J. M. Małecki, M.-F. Odonohue, Y. Kim, M. E. Jakobsson, L. Gessa, R. Pinto, J. Wu, E. Davydova, A. Moen, J. V Olsen, B. Thiede, P.-E. Gleizes, S. A. Leidel, P. Ø. Falnes, Human METTL18 is a histidine-specific methyltransferase that targets RPL3 and affects ribosome biogenesis and function. Nucleic Acids Research. 49, 3185–3203 (2021).

5. Q. Al-Hadid, K. Roy, W. Munroe, M. C. Dzialo, G. F. Chanfreau, S. G. Clarke, Histidine Methylation of Yeast Ribosomal Protein Rpl3p Is Required for Proper 60S Subunit Assembly. Molecular and Cellular Biology. 34, 2903–2916 (2014).

6. M. E. Jakobsson, Enzymology and significance of protein histidine methylation. Journal of Biological Chemistry. 297, 101130 (2021).

7. R. J. Quinlan, M. D. Sweeney, L. Lo Leggio, H. Otten, J.-C. N. Poulsen, K. S. Johansen, K. B. R. M. Krogh, C. I. Jørgensen, M. Tovborg, A. Anthonsen, T. Tryfona, C. P. Walter, P. Dupree, F. Xu, G. J. Davies, P. H. Walton, Insights into the oxidative degradation of cellulose by a copper metalloenzyme that exploits biomass components. Proc. Natl. Acad. Sci. U.S.A. 108, 15079–15084 (2011).

8. T. Tandrup, K. E. H. Frandsen, K. S. Johansen, J. G. Berrin, L. Lo Leggio, Recent insights into lytic polysaccharide monooxygenases (LPMOs). Biochemical Society Transactions. 46, 1431–1447 (2018).

9. J. Ipsen, M. Hallas-Møller, S. Brander, L. Lo Leggio, K. S. Johansen, Lytic polysaccharide monooxygenases and other histidine-brace copper proteins: Structure, oxygen activation and biotechnological applications. Biochemical Society Transactions. 49, 531–540 (2021).

10. K. S. Johansen, Discovery and industrial applications of lytic polysaccharide mono-oxygenases. Biochemical Society Transactions. 44, 143–149 (2016).

11. J. S. M. Loose, Z. Forsberg, M. W. Fraaije, V. G. H. Eijsink, G. Vaaje-kolstad, A rapid quantitative activity assay shows that the Vibrio cholerae colonization factor GbpA is an active lytic polysaccharide monooxygenase. FEBS Letters. 588, 3435–3440 (2014).

12. F. Askarian, S. Uchiyama, H. Masson, H. V. Sørensen, O. Golten, A. C. Bunæs, S. Mekasha, Å. K. Røhr, E. Kommedal, J. A. Ludviksen, M. Ø. Arntzen, B. Schmidt, R. H. Zurich, N. M. van Sorge, V. G. H. Eijsink, U. Krengel, T. E. Mollnes, N. E. Lewis, V. Nizet, G. Vaaje-Kolstad, The lytic polysaccharide monooxygenase CbpD promotes Pseudomonas aeruginosa virulence in systemic infection. Nature Communications. 12, 1230 (2021).

13. Á. Polonio, D. Fernández-Ortuño, A. Vicente, A. Pérez-García, A haustorial-expressed lytic polysaccharide monooxygenase from the cucurbit powdery mildew pathogen Podosphaera xanthii contributes to the suppression of chitin-triggered immunity. Molecular Plant Pathology. 22, 580–601 (2021).

14. D. M. Petrović, B. Bissaro, P. Chylenski, M. Skaugen, M. Sørlie, M. S. Jensen, F. L. Aachmann, G. Courtade, A. Várnai, V. G. H. Eijsink, Methylation of the N-terminal histidine protects a lytic polysaccharide monooxygenase from auto-oxidative inactivation: Role of Histidine Methylation in Fungal LPMOs. Protein Science. 27, 1636–1650 (2018).

15. S. Banerjee, S. J. Muderspach, T. Tandrup, K. E. H. Frandsen, R. K. Singh, J. Ø. Ipsen, C. Hernández-Rollán, M. H. H. Nørholm, M. J. Bjerrum, K. S. Johansen, L. Lo Leggio, Protonation State of an Important Histidine from High Resolution Structures of Lytic Polysaccharide Monooxygenases. Biomolecules. 12, 194 (2022).

16. L. A. Kelley, S. Mezulis, C. M. Yates, M. N. Wass, M. J. E. Sternberg, The Phyre2 web portal for protein modeling, prediction and analysis. Nat Protoc. 10, 845–858 (2015).

17. M. Blum, H.-Y. Chang, S. Chuguransky, T. Grego, S. Kandasaamy, A. Mitchell, G. Nuka, T. Paysan-Lafosse, M. Qureshi, S. Raj, L. Richardson, G. A. Salazar, L. Williams, P. Bork, A. Bridge, J. Gough, D. H. Haft, I. Letunic, A. Marchler-Bauer, H. Mi, D. A. Natale, M. Necci, C. A. Orengo, A. P. Pandurangan, C. Rivoire, C. J. A. Sigrist, I. Sillitoe, N. Thanki, P. D. Thomas, S. C. E. Tosatto, C. H. Wu, A. Bateman, R. D. Finn, The InterPro protein families and domains database: 20 years on. Nucleic Acids Research. 49, D344–D354 (2021).

18. J. Mistry, S. Chuguransky, L. Williams, M. Qureshi, G. A. Salazar, E. L. L. Sonnhammer, S. C. E. Tosatto, L. Paladin, S. Raj, L. J. Richardson, R. D. Finn, A. Bateman, Pfam: The protein families database in 2021. Nucleic Acids Research. 49, D412–D419 (2021).

19. M. Haddad Momeni, M. L. Leth, C. Sternberg, E. Schoof, M. W. Nielsen, J. Holck, C. T. Workman, J. B. Hoof, M. Abou Hachem, Loss of AA13 LPMOs impairs degradation of resistant starch and reduces the growth of Aspergillus nidulans. Biotechnol Biofuels. 13, 135 (2020).

20. C. J. A. Sigrist, E. de Castro, L. Cerutti, B. A. Cuche, N. Hulo, A. Bridge, L. Bougueleret, Xenarios, New and continuing developments at PROSITE. Nucleic Acids Research. 41, D344–D347 (2012).

21. C. S. Nødvig, J. B. Nielsen, M. E. Kogle, U. H. Mortensen, A CRISPR-Cas9 System for Genetic Engineering of Filamentous Fungi. PLOS ONE. 10, e0133085 (2015).

22. C. S. Nødvig, J. B. Hoof, M. E. Kogle, Z. D. Jarczynska, J. Lehmbeck, D. K. Klitgaard, U. H. Mortensen, Efficient oligo nucleotide mediated CRISPR-Cas9 gene editing in Aspergilli. Fungal Genetics and Biology. 115, 78–89 (2018).

23. L. J. McGuffin, K. Bryson, D. T. Jones, The PSIPRED protein structure prediction server. Bioinformatics. 16, 404–405 (2000).

24. T. Nugent, D. T. Jones, Transmembrane protein topology prediction using support vector machines. BMC Bioinformatics. 10, 159 (2009).

25. M. E. Jakobsson, J. M. Małecki, L. Halabelian, B. S. Nilges, R. Pinto, S. Kudithipudi, S. Munk, E. Davydova, F. R. Zuhairi, C. H. Arrowsmith, A. Jeltsch, S. A. Leidel, J. V. Olsen, P. Ø. Falnes, The dual methyltransferase METTL13 targets N terminus and Lys55 of eEF1A and modulates codon-specific translation rates. Nat Commun. 9, 3411 (2018).

26. A. W. Struck, M. L. Thompson, L. S. Wong, J. Micklefield, S-Adenosyl-Methionine-Dependent Methyltransferases: Highly Versatile Enzymes in Biocatalysis, Biosynthesis and Other Biotechnological Applications. ChemBioChem. 13, 2642–2655 (2012).

27. J. R. Wortman, J. M. Gilsenan, V. Joardar, J. Deegan, J. Clutterbuck, M. R. Andersen, D. Archer, M. Bencina, G. Braus, P. Coutinho, H. von Döhren, J. Doonan, A. J. M. Driessen, P. Durek, E. Espeso, E. Fekete, M. Flipphi, C. G. Estrada, S. Geysens, G. Goldman, P. W. J. de Groot, K. Hansen, S. D. Harris, T. Heinekamp, K. Helmstaedt, B. Henrissat, G. Hofmann, T. Homan, T. Horio, H. Horiuchi, S. James, M. Jones, L. Karaffa, Z. Karányi, M. Kato, N. Keller, D. E. Kelly, J. A. K. W. Kiel, J.-M. Kim, I. J. van der Klei, F. M. Klis, A. Kovalchuk, N. Krasevec, C. P. Kubicek, B. Liu, A. Maccabe, V. Meyer, P. Mirabito, M. Miskei, M. Mos, J. Mullins, D. R. Nelson, J. Nielsen, B. R. Oakley, S. A. Osmani, T. Pakula, A. Paszewski, I. Paulsen, S. Pilsyk, I. Pócsi, P. J. Punt, A. F. J. Ram, Q. Ren, X. Robellet, G. Robson, B. Seiboth, P. van Solingen, T. Specht, J. Sun, N. Taheri-Talesh, N. Takeshita, D. Ussery, P. A. vanKuyk, H. Visser, P. J. I. van de Vondervoort, R. P. de Vries, J. Walton, X. Xiang, Y. Xiong, A. P. Zeng, B. W. Brandt, M. J. Cornell, C. A. M. J. J. van den Hondel, J. Visser, S. G. Oliver, G. Turner, The 2008 update of the Aspergillus nidulans genome annotation: a community effort. Fungal Genet Biol. 46 Suppl 1, S2–13 (2009).

28. H. Wu, J. Min, Y. Ikeguchi, H. Zeng, A. Dong, P. Loppnau, A. E. Pegg, A. N. Plotnikov, Structure and Mechanism of Spermidine Synthases. Biochemistry. 46, 8331–8339 (2007).

29. Y. Ikeguchi, M. C. Bewley, A. E. Pegg, Aminopropyltransferases: Function, Structure and Genetics. The Journal of Biochemistry. 139, 1–9 (2006).

30. Y. Amano, I. Namatame, Y. Tateishi, K. Honboh, E. Tanabe, T. Niimi, H. Sakashita, Structural insights into the novel inhibition mechanism of Trypanosoma cruzi spermidine synthase. Acta Cryst D. 71, 1879–1889 (2015).

31. R. P. D. Bank, RCSB PDB - 1IY9: Crystal structure of spermidine synthase, (available at https://www.rcsb.org/structure/1iy9).

32. I. Letunic, P. Bork, Interactive Tree Of Life (iTOL) v5: an online tool for phylogenetic tree display and annotation. Nucleic Acids Research. 49, W293–W296 (2021).

33. L. Holm, Dali server: structural unification of protein families. Nucleic Acids Research. 50, W210–W215 (2022).

34. R. W. Sabnis, T. G. Deligeorgiev, M. N. Jachak, T. S. Dalvi, DiOC _6_ (3): a Useful Dye for Staining the Endoplasmic Reticulum. Biotechnic & Histochemistry. 72, 253–258 (1997).

35. D. C. Anyaogu, A. H. Hansen, J. B. Hoof, N. I. Majewska, F. J. Contesini, J. T. Paul, K. F. Nielsen, T. J. Hobley, S. Yang, H. Zhang, M. Betenbaugh, U. H. Mortensen, Glycoengineering of Aspergillus nidulans to produce precursors for humanized N-glycan structures. Metab Eng. 67, 153–163 (2021).

36. C. Hernández-Rollán, K. B. Falkenberg, M. Rennig, A. B. Bertelsen, J. Ø. Ipsen, S. Brander, D. O. Daley, K. S. Johansen, M. H. H. Nørholm, LyGo: A Platform for Rapid Screening of Lytic Polysaccharide Monooxygenase Production. ACS Synthetic Biology. 10, 897–906 (2021).

37. L. Rieder, K. Ebner, A. Glieder, M. Sørlie, Novel molecular biological tools for the efficient expression of fungal lytic polysaccharide monooxygenases in Pichia pastoris. Biotechnology for Biofuels. 14, 1–17 (2021).

38. M. Tanghe, B. Danneels, A. Camattari, A. Glieder, I. Vandenberghe, B. Devreese, I. Stals, T. Desmet, Recombinant Expression of Trichoderma reesei Cel61A in Pichia pastoris: Optimizing Yield and N-terminal Processing. Molecular Biotechnology. 57, 1010–1017 (2015).

39. D. R. Brademan, N. M. Riley, N. W. Kwiecien, J. J. Coon, Interactive Peptide Spectral Annotator: A Versatile Web-based Tool for Proteomic Applications. Molecular & Cellular Proteomics. 18, S193–S201 (2019).

40. P. S. Pathinayake, A. C.-Y. Hsu, P. A. B. Wark, PAT in the ER for Transmembrane Protein Folding. Trends in Biochemical Sciences. 45, 1007–1008 (2020).

41. J. J. Kattla, W. B. Struwe, M. Doherty, B. Adamczyk, R. Saldova, P. M. Rudd, M. P. Campbell, Protein Glycosylation (Elsevier B.V., Second Edi., 2011), vol. 3.

42. G. Walsh, R. Jefferis, Post-translational modifications in the context of therapeutic proteins. Nature Biotechnology. 24, 1241–1252 (2006).

43. S. G. W. Kaminskyj, Fundamentals of growth, storage, genetics and microscopy of Aspergillus nidulans. Fungal Genetics Reports. 48, 25–31 (2001).

44. D. J. Cove, The induction and repression of nitrate reductase in the fungus Aspergillus nidulans. Biochimica et Biophysica Acta (BBA) - Enzymology and Biological Oxidation. 113, 51–56 (1966).

45. H. H. Nour-Eldin, F. Geu-Flores, B. A. Halkier, “USER Cloning and USER Fusion: The Ideal Cloning Techniques for Small and Big Laboratories” in Plant Secondary Metabolism Engineering: Methods and Applications, A. G. Fett-Neto, Ed. (Humana Press, Totowa, NJ, 2010; https://doi.org/10.1007/978-1-60761-723-5_13), Methods in Molecular Biology, pp. 185–200.

46. A. M. Cavaleiro, S. H. Kim, S. Seppälä, M. T. Nielsen, M. H. H. Nørholm, Accurate DNA Assembly and Genome Engineering with Optimized Uracil Excision Cloning. ACS Synth. Biol. 4, 1042–1046 (2015).

47. B. G. Hansen, B. Salomonsen, M. T. Nielsen, J. B. Nielsen, N. B. Hansen, K. F. Nielsen, T. B. Regueira, J. Nielsen, K. R. Patil, U. H. Mortensen, Versatile Enzyme Expression and Characterization System for Aspergillus nidulans, with the Penicillium brevicompactum Polyketide Synthase Gene from the Mycophenolic Acid Gene Cluster as a Test Case▿. Appl Environ Microbiol. 77, 3044–3051 (2011).

48. F. Teufel, J. J. Almagro Armenteros, A. R. Johansen, M. H. Gíslason, S. I. Pihl, K. D. Tsirigos, O. Winther, S. Brunak, G. von Heijne, H. Nielsen, SignalP 6.0 predicts all five types of signal peptides using protein language models. Nat Biotechnol, 1–3 (2022).

49. C. C. Lee, T. G. Williams, D. W. S. Wong, G. H. Robertson, An episomal expression vector for screening mutant gene libraries in Pichia pastoris. Plasmid. 54, 80–85 (2005).

50. M. L. Nielsen, L. Albertsen, G. Lettier, J. B. Nielsen, U. H. Mortensen, Efficient PCR-based gene targeting with a recyclable marker for Aspergillus nidulans. Fungal Genetics and Biology. 43, 54–64 (2006).

51. J. R. Wiśniewski, F. Z. Gaugaz, Fast and Sensitive Total Protein and Peptide Assays for Proteomic Analysis. Anal. Chem. 87, 4110–4116 (2015).

52. T. S. Batth, MaximA. X. Tollenaere, P. Rüther, A. Gonzalez-Franquesa, B. S. Prabhakar, S. Bekker-Jensen, A. S. Deshmukh, J. V. Olsen, Protein Aggregation Capture on Microparticles Enables Multipurpose Proteomics Sample Preparation*. Molecular & Cellular Proteomics. 18, 1027–1035 (2019).

53. A. Lundby, J. V. Olsen, “GeLCMS for In-Depth Protein Characterization and Advanced Analysis of Proteomes” in Gel-Free Proteomics: Methods and Protocols, K. Gevaert, J. Vandekerckhove, Eds. (Humana Press, Totowa, NJ, 2011; https://doi.org/10.1007/978-1-61779-148-2_10), Methods in Molecular Biology, pp. 143–155.

54. J. Cox, M. Mann, MaxQuant enables high peptide identification rates, individualized p.p.b.-range mass accuracies and proteome-wide protein quantification. Nat Biotech. 26, 1367–1372 (2008).

55. S. Kapell, M. E. Jakobsson, Large-scale identification of protein histidine methylation in human cells. NAR Genomics and Bioinformatics. 3, nqab045 (2021).

56. B. MacLean, D. M. Tomazela, N. Shulman, M. Chambers, G. L. Finney, B. Frewen, R. Kern, D. L. Tabb, D. C. Liebler, M. J. MacCoss, Skyline: an open source document editor for creating and analyzing targeted proteomics experiments. Bioinformatics. 26, 966–968 (2010).

57. S. Tyanova, T. Temu, P. Sinitcyn, A. Carlson, M. Y. Hein, T. Geiger, M. Mann, J. Cox, The Perseus computational platform for comprehensive analysis of (prote)omics data. Nat Methods. 13, 731–740 (2016).

