## Supplementary Information for "A membrane integral methyltransferase catalysing N-terminal histidine methylation of lytic polysaccharide monooxygenases"

*Shared first-authors

#Corresponding authors

### Mass spectrometry data

#### Supplementary Figure S1 – Gene knockout PRM assay

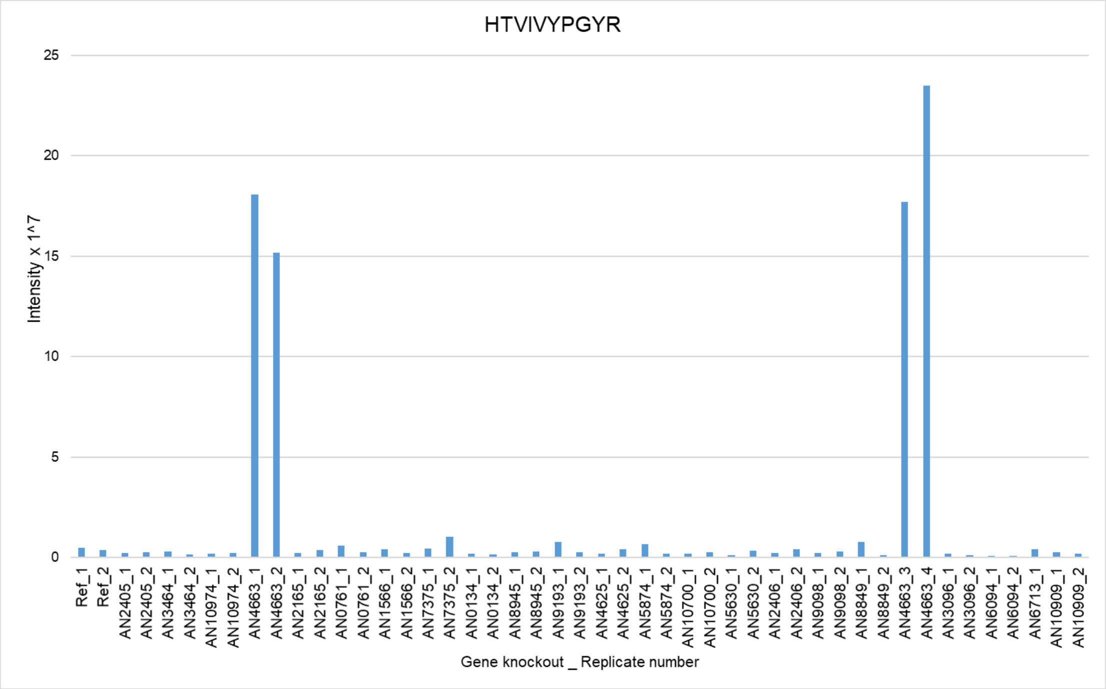

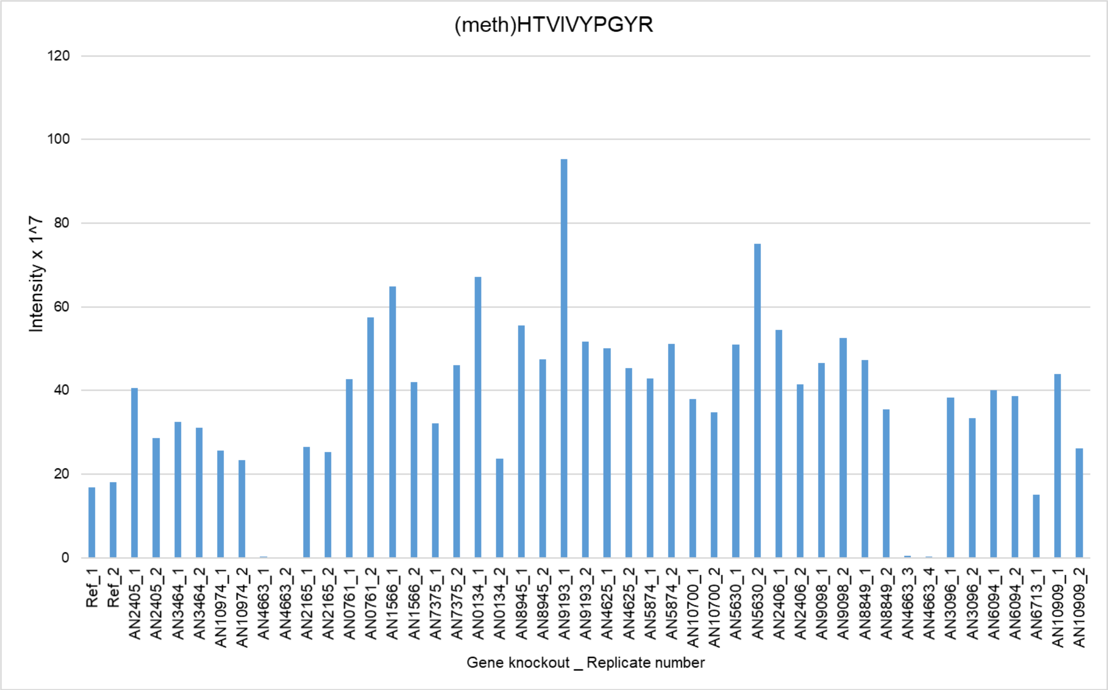

**Supplementary Figure S1 – Gene knockout PRM assay.** PRM quantification of Q5B428 N-terminal histidine methylated (609.8351 m/z) and unmethylated (602.8273 m/z) HTVIVYPGYR peptide for the different gene knockouts shown above with the two biological replicates. Additional biological replicates for AN4663 knockout were performed to confirm the results.

### *In silico* analysis

#### Supplementary Figure S2 – Secondary structure prediction with PSIPRED and MEMSAT

A

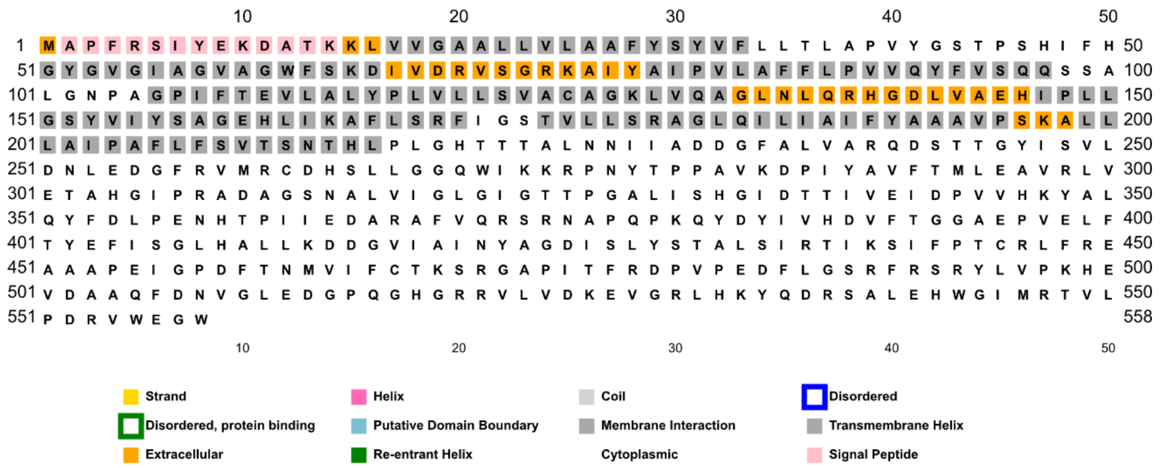

B

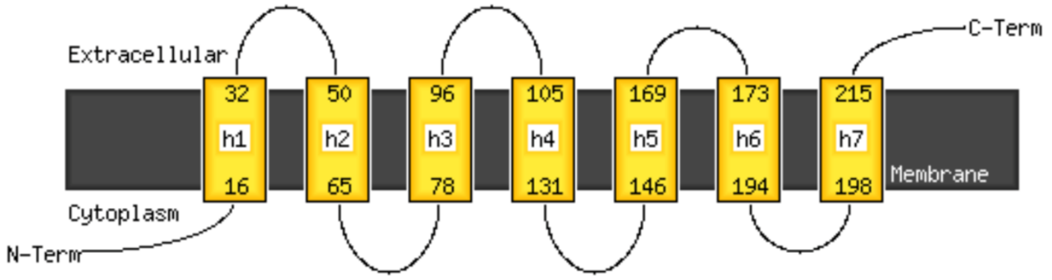

**Supplementary Figure S2 – Secondary structure prediction with PSIPRED and MEMSAT.** Secondary structure was prediction of the AN4663 protein sequence analyzed using A) PSIPRED shows the predicted secondary structure regions and B) MEMSAT analysis for prediction of transmembrane protein topology of AN4663.

#### Supplementary Figure S3 – *In silico* analysis of the soluble domain

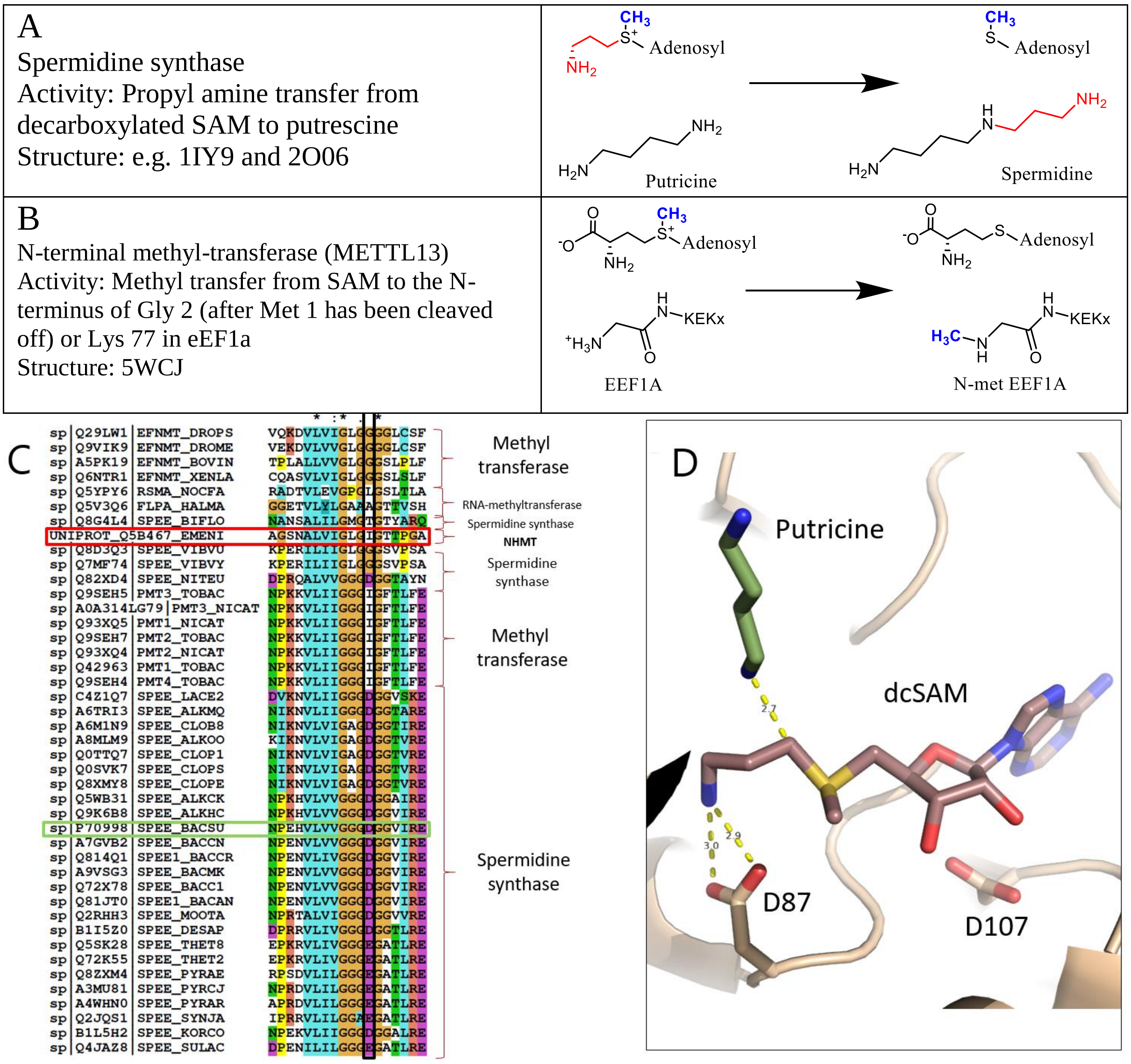

**Supplementary Figure S3 – *In silico* analysis of the soluble domain.** The substrate and co-substrates differ between **A)** spermidine synthases and **B)** methyl transferases. **C)** BLAST search of the soluble domain of NHMT (residues 216-558) against the manually curated database uniprotkb_swissprot, using the BLOSUM-45 matrix and an exp cutoff at 1.0 C). Framed in red, NHMT. Framed in green, the *B. subtilis* sequence that contains the spermidine synthase motif GxG(DE)G and is the most homologue sequence to NHMT where the experimental three-dimensional structure is available. The discriminating feature of spermidine synthases is the glycine rich loop where the acidic residue binds to the amine of decarboxylated SAM. **D)** The importance of the acidic residue is shown in an amalgam of structures from human (pdb:2O06), *T. cruzi* (pdb:4YV0) and *B. subtilis* (pdb:1IY9) spermidine synthases where the putrescine aligns perfectly for a nucleophilic attack on the decarboxylated SAM propylene moiety.

#### Supplementary Figure S4 – Sequence analysis of the 7TM domain shows that it is unique

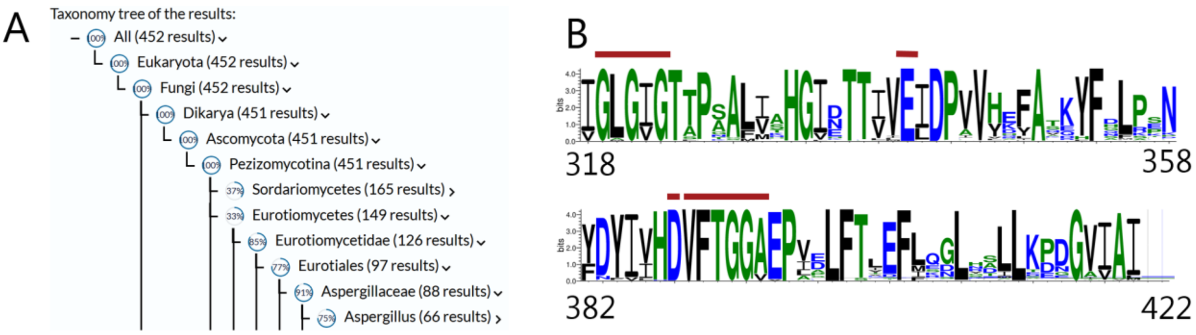

C

| # No | Chain | Z | rmsd | %id | Description |
|---|---|---|---|---|---|
| 1 | 6rw3-B | 10.7 | 4.2 | 8 | HEXOSE TRANSPORTER 1; |
| 2 | 6kkk-A | 10.3 | 4.3 | 9 | SUGAR EFFLUX TRANSPORTER; |
| 3 | 6g9x-A | 10.3 | 3.8 | 10 | MAJOR FACILITATOR SUPERFAMILY MFS_1; |
| 4 | 4zp0-A | 10.2 | 4.1 | 9 | MULTIDRUG TRANSPORTER MDFA; |
| 5 | 7bc6-A | 9.9 | 4.1 | 11 | PROTON-COUPLED FOLATE TRANSPORTER; |
| 6 | 6t1z-A | 9.7 | 4.5 | 9 | LMRP INTEGRAL MEMBRANE PROTEIN; |
| 7 | 7nqk-A | 9.6 | 3.9 | 7 | SOLUTE CARRIER FAMILY 15 MEMBER 2; |
| 8 | 6s7v-A | 9.5 | 4.1 | 13 | MFS TRANSPORTER; |
| 9 | 4u4t-A | 9.3 | 4.3 | 11 | NITRATE/NITRITE TRANSPORTER NARK; |
| 10 | 2gfp-A | 9.2 | 3.8 | 8 | MULTIDRUG RESISTANCE PROTEIN D; |
| 11 | 7d5p-B | 9.2 | 3.9 | 9 | DRUG TRANSPORTER, PUTATIVE; |
| 12 | 7c76-B | 9.1 | 4.6 | 7 | TOLL-LIKE RECEPTOR 3; |
| 13 | 6s4m-A | 9.0 | 3.8 | 12 | MAJOR FACILITATOR SUPERFAMILY .. |
| 14 | 6gz9-A | 9.0 | 4.2 | 12 | PEPTIDE ABC TRANSPORTER PERMEASE; |
| 15 | 6e9c-A | 8.7 | 4.1 | 7 | MAJOR FACILITATOR FAMILY TRANSPORTER; |
| 16 | 3wdo-A | 8.6 | 4.0 | 12 | MFS TRANSPORTER; |
| 17 | 6v4d-A | 8.5 | 4.2 | 9 | VESICULAR GLUTAMATE TRANSPORTER 2; |
| 18 | 6e9n-A | 8.4 | 4.0 | 8 | D-GALACTONATE TRANSPORT; |
| 19 | 1pw4-A | 8.4 | 4.0 | 8 | GLYCEROL-3-PHOSPHATE TRANSPORTER; |
| 20 | 6wbv-A | 8.0 | 4.3 | 9 | HEPCIDIN; |
| 21 | 3o7q-A | 7.9 | 3.7 | 12 | L-FUCOSE-PROTON SYMPORTER; |
| 22 | 7d19-A | 7.8 | 4.0 | 8 | NUCLEOSIDE PERMEASE NUPG; |

**Supplementary Figure S4 – Sequence analysis of the 7TM domain shows that it is unique. A)** 452 sequences of homologous 7TM domains. All but one found in the Pezizomycotina phylum (filamentous ascomycetes). The sequence of the 7TM domain (residues 1-215) was used as input for a pBLAST search against the uniprotkb_refprotswissprot database using the BLOSUM-45 matrix and an exp cutoff at 1e-4. **B)** Alignment of the soluble domain linked to the 452 transmembrane domain sequences by MUSCLE v. 3.8 and visualized as sequence motifs in WebLogo 3.0. The region 318-358 shows the methyl transferase motif at 319-323 as well as the SAM binding glutamate in position 340. The highly conserved segment 389-394 is uniquely found in proteins with the nHMT 7TM domain. **C)** The first 22 hits found by search in the Dali25 database with the AlphaFold2 model of the An-nHMT 7TM domain. The structures of these proteins were investigated manually and none of them had 7 helices.

### Protein expression

#### Supplementary Figure S5 – LsAA9A secretion with the α-MF signal peptide

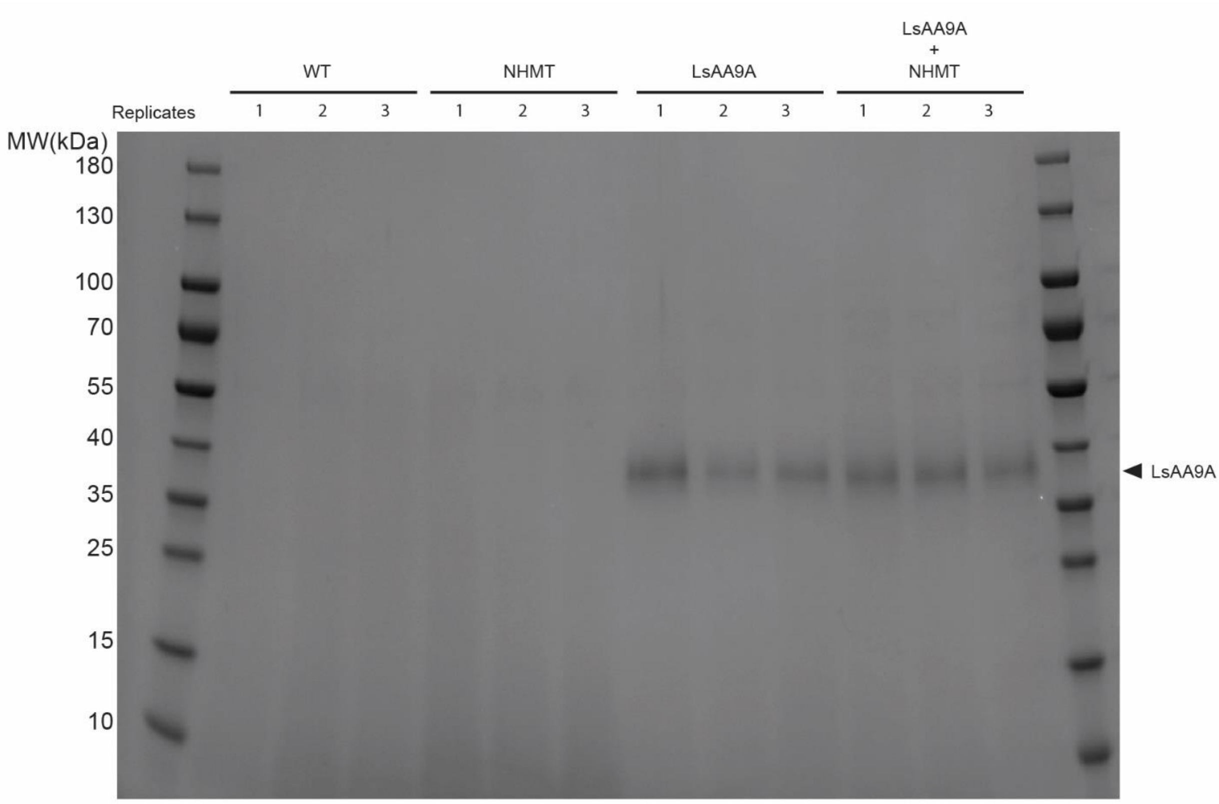

**Supplementary Figure S5 – LsAA9A secretion with the α-MF signal peptide.** SDS-PAGE of secreted samples into the supernatant from the co-expression of LsAA9A and the methyltransferase AN4663 when the LsAA9A is expressed using the α-MF signal peptide in biological triplicates. WT (wild-type, parent strain GS115), NHMT (N-terminal histidine methyltransferase expressed from an episomal plasmid), LsAA9A (LsAA9A secreted into the supernatant), LsAA9A + NHMT (LsAA9A co-expressed with the N-terminal histidine methyltransferase). MW (molecular weight in kDa ladder). LsAA9A expected molecular weight without glycosylation is 25.2 kDa. The arrow indicates were the secreted LsAA9A is expected.

#### Supplementary Figure S6 – LsAA9A secretion with the Amy signal peptide

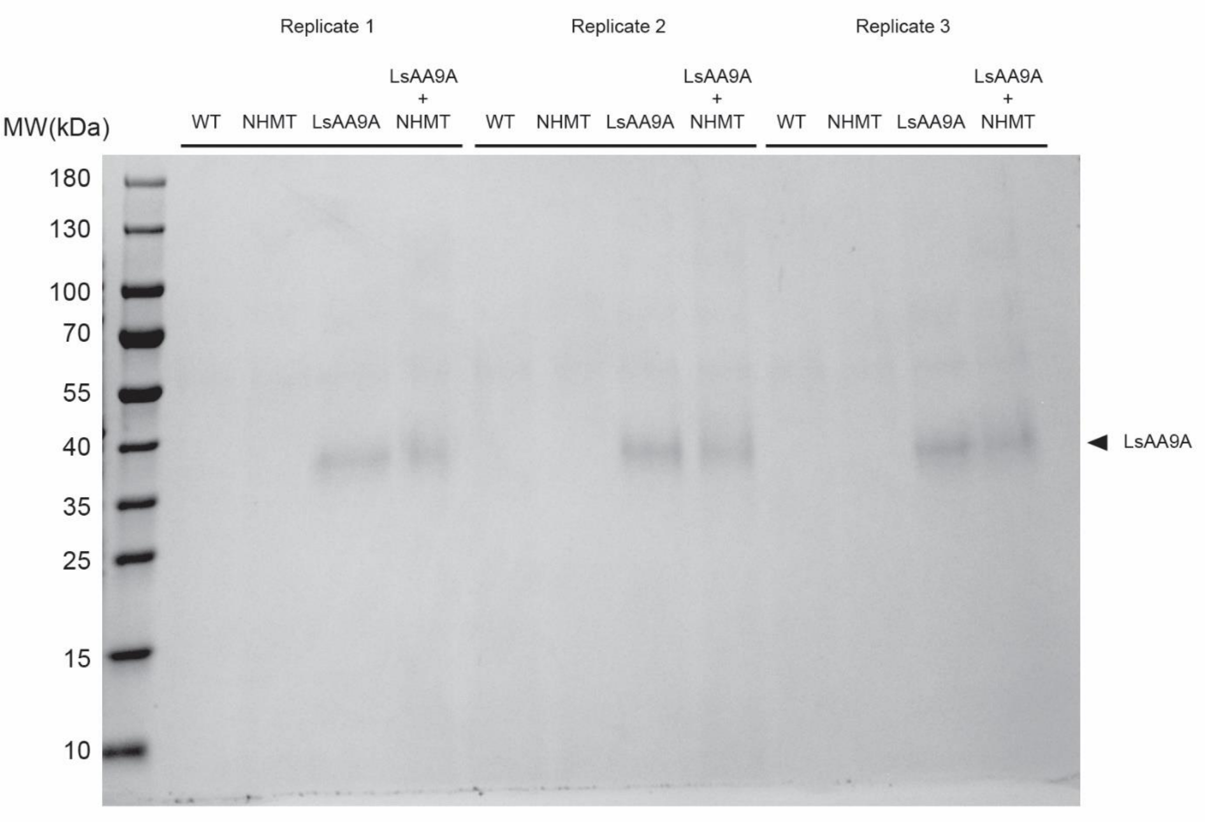

**Supplementary Figure S6 – LsAA9A secretion with the Amy signal peptide.** SDS-PAGE of secreted samples into the supernatant from the co-expression of LsAA9A and the methyltransferase AN4663 when the LsAA9A is expressed using the Amy signal peptide in biological triplicates. WT (wild-type, parent strain GS115), NHMT (N-terminal histidine methyltransferase expressed from an episomal plasmid), LsAA9A (LsAA9A secreted into the supernatant), LsAA9A + NHMT (LsAA9A co-expressed with the N-terminal histidine methyltransferase). MW (molecular weight in kDa ladder). LsAA9A expected molecular weight without glycosylation is 25.2 kDa. The arrow indicates were the secreted LsAA9A is expected.

#### Supplementary Figure S7 – LsAA9A secretion with its native signal peptide

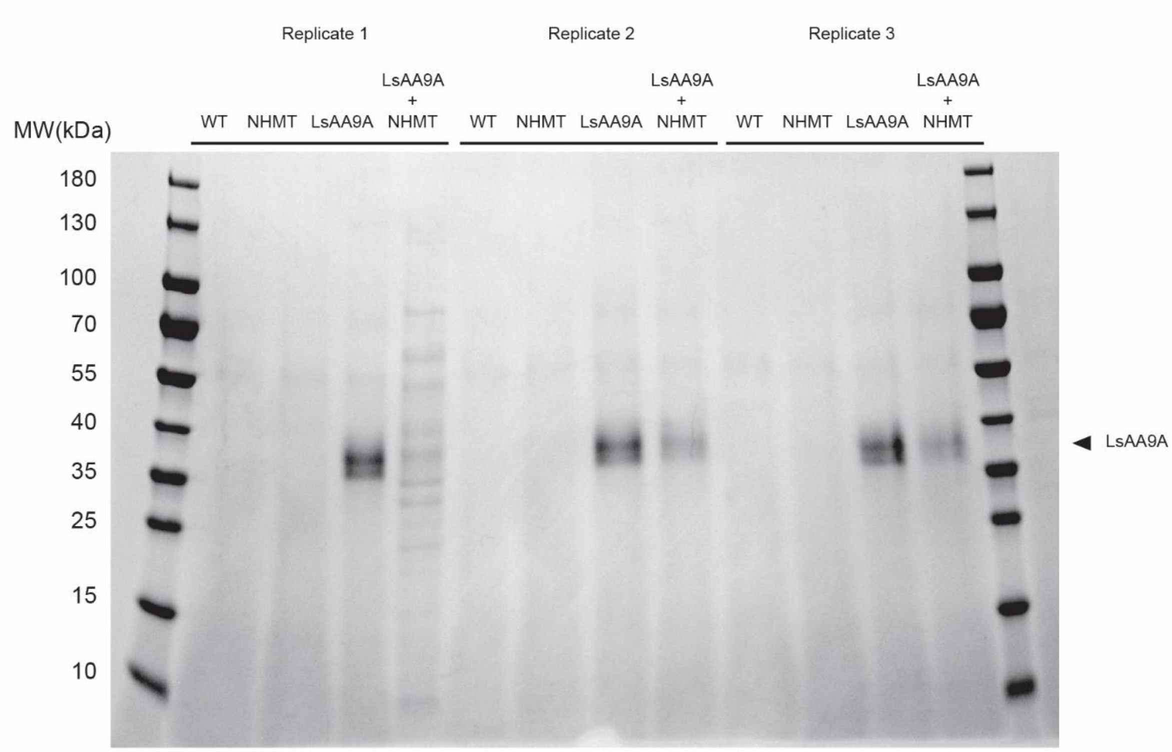

**Supplementary Figure S7 – LsAA9A secretion with its native signal peptide**. SDS-PAGE of secreted samples into the supernatant from the co-expression of LsAA9A and the methyltransferase AN4663 when the LsAA9A is expressed using its own native signal peptide in biological triplicates. WT (wild-type, parent strain GS115), NHMT (N-terminal histidine methyltransferase expressed from an episomal plasmid), LsAA9A (LsAA9A secreted into the supernatant), LsAA9A + NHMT (LsAA9A co-expressed with the N-terminal histidine methyltransferase). MW (molecular weight in kDa ladder). LsAA9A expected molecular weight without glycosylation is 25.2 kDa. The arrow indicates were the secreted LsAA9A is expected.

#### Supplementary Figure S8 – LsAA9A processing with alpha-mating factor signal peptide

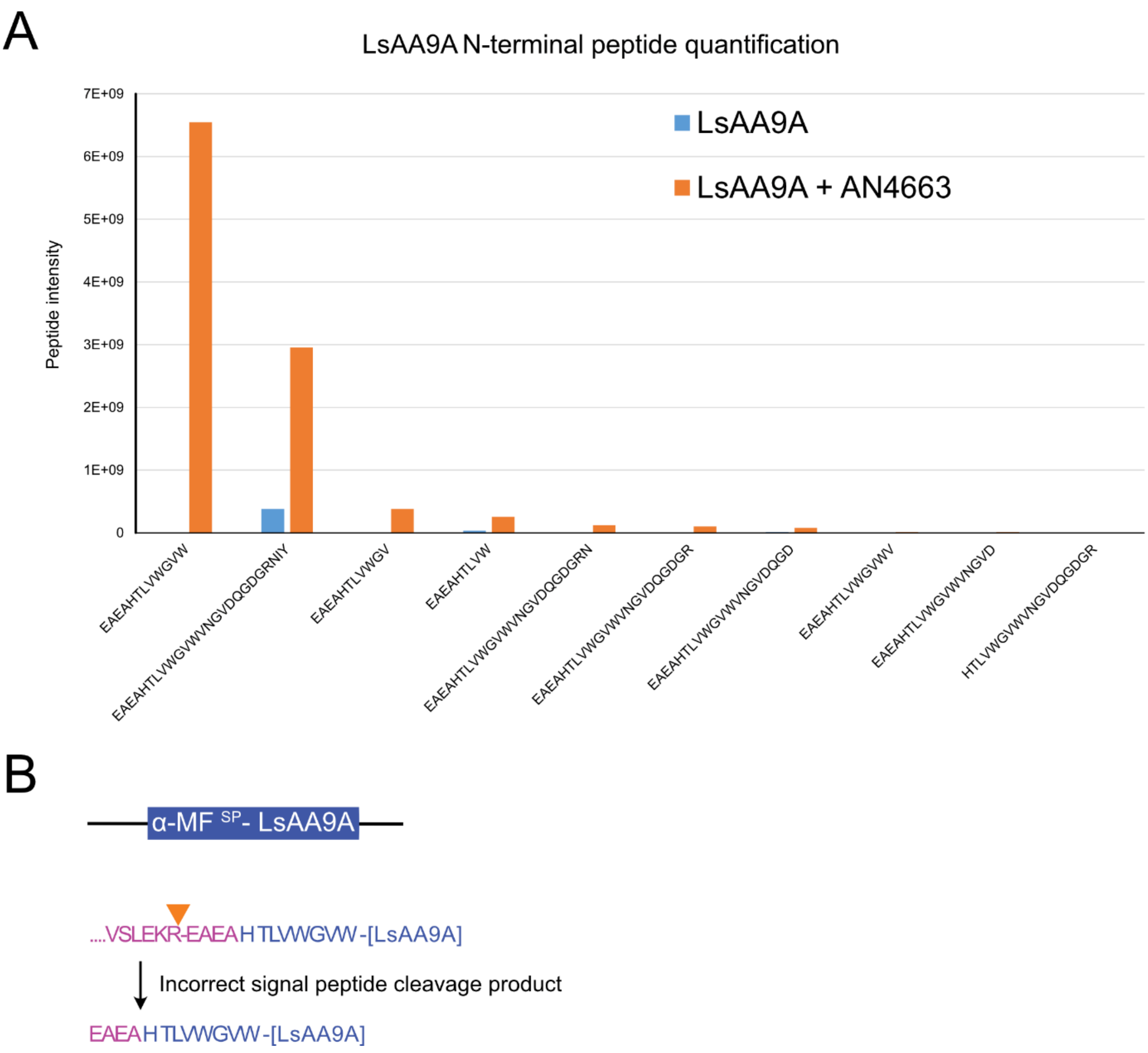

**Supplementary Figure S8 – LsAA9A processing with alpha-mating factor signal peptide.** LsAA9A N-terminal peptide MS identification and quantification with alpha-mating factor secretion signal peptide fused to the N-terminal of mature LsAA9A protein sequence. A) Quantification of identified N-terminal peptides of strains expressing LsAA9A (blue) with a alpha-mating factor signal peptide co-expressed with AN4663. B) Illustration of mature LsAA9A sequence with the addition of alpha-mating factor signal peptide.

#### Supplementary Figure S9 – Tf10A secretion with the Amy signal peptide

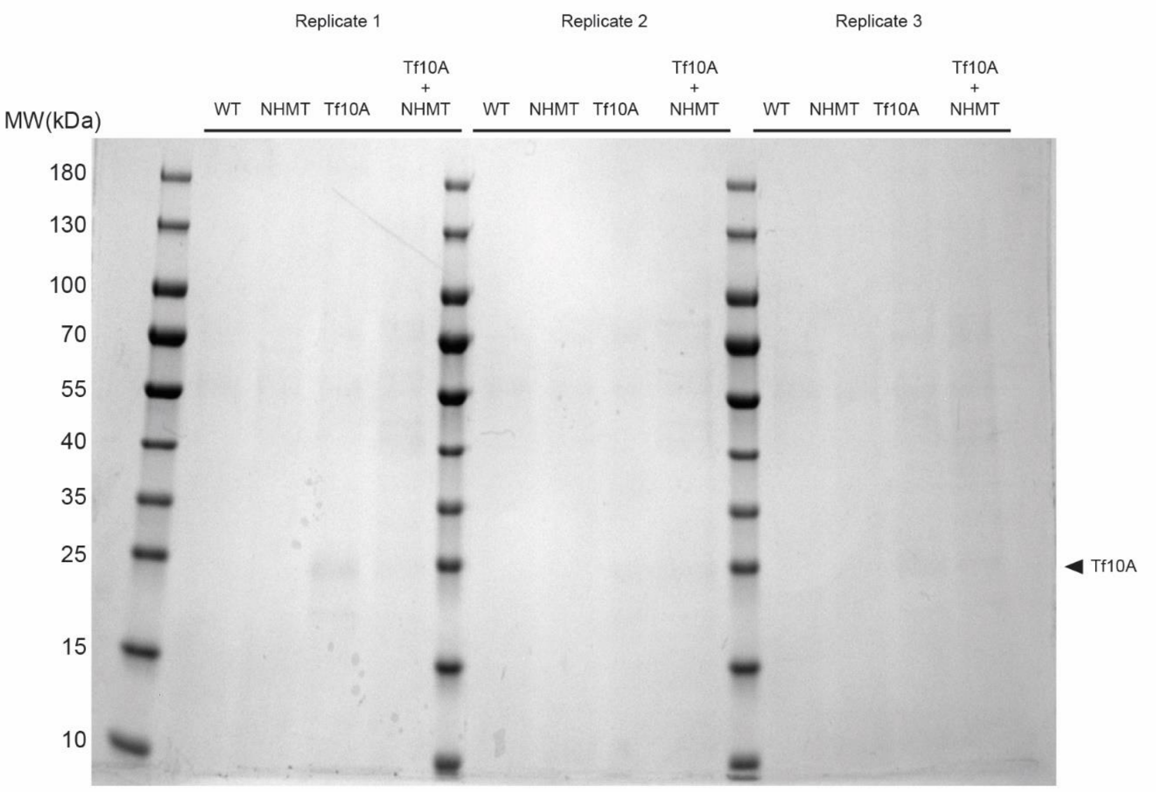

**Supplementary Figure S9 – Tf10A secretion with the Amy signal peptide**. SDS-PAGE of secreted samples into the supernatant from the co-expression of Tf10A and the methyltransferase AN4663 when the Tf10A is expressed using the Amy signal peptide in biological triplicates. WT (wild-type, parent strain GS115), NHMT (N-terminal histidine methyltransferase expressed from an episomal plasmid), Tf10A (Tf10A secreted into the supernatant), Tf10A + NHMT (Tf10A co-expressed with the N-terminal histidine methyltransferase). MW (molecular weight in kDa ladder). Tf10A expected molecular weight without glycosylation is 21.3 kDa. The arrow indicates were the secreted Tf10A is expected.

Table S1 – C8V530 pfam predication

| Source | Domain | Start | End | Gathering threshold (bits) | | Score (bits) | | E-value | |
|---|---|---|---|---|---|---|---|---|---|
| | | | | Sequence | Domain | Sequence | Domain | Sequence | Domain |
| **sig_p** | n/a | 1 | 22 | n/a | n/a | n/a | n/a | n/a | n/a |
| **low_complexity** | n/a | 6 | 21 | n/a | n/a | n/a | n/a | n/a | n/a |
| **Pfam** | AA9 | 23 | 242 | 27.8 | 27.8 | 226 | 225.7 | 1.50E-63 | 1.90E-63 |
| **disorder** | n/a | 50 | 53 | n/a | n/a | n/a | n/a | n/a | n/a |
| **disorder** | n/a | 58 | 59 | n/a | n/a | n/a | n/a | n/a | n/a |
| **disorder** | n/a | 63 | 64 | n/a | n/a | n/a | n/a | n/a | n/a |
| **disorder** | n/a | 76 | 79 | n/a | n/a | n/a | n/a | n/a | n/a |
| **disorder** | n/a | 216 | 221 | n/a | n/a | n/a | n/a | n/a | n/a |
| **disorder** | n/a | 267 | 271 | n/a | n/a | n/a | n/a | n/a | n/a |

**Table S1.** This table includes pfam predicted domain of the *A. nidulans* protein C8V530 (Uniprot identifier). An auxiliary activity family 9 (AA9) domain with pfam accession PF03443 is predicted within C8V530.

### Table S2 - Q5B1W7 pfam predication

| Source | Domain | Start | End | Gathering threshold (bits) | | Score (bits) | | E-value | |
|---|---|---|---|---|---|---|---|---|---|
| | | | | Sequence | Domain | Sequence | Domain | Sequence | Domain |
| **sig_p** | n/a | 1 | 15 | n/a | n/a | n/a | n/a | n/a | n/a |
| **Pfam** | LPMO_10 | 19 | 248 | 23 | 23 | 28.1 | 26.9 | 0.0092 | 0.022 |
| **low_complexity** | n/a | 252 | 281 | n/a | n/a | n/a | n/a | n/a | n/a |
| **disorder** | n/a | 259 | 270 | n/a | n/a | n/a | n/a | n/a | n/a |
| **Pfam** | CBM_20 | 283 | 379 | 23.3 | 23.3 | 121 | 120.2 | 4.80E-32 | 8.40E-32 |
| **low_complexity** | n/a | 315 | 329 | n/a | n/a | n/a | n/a | n/a | n/a |
| **disorder** | n/a | 360 | 361 | n/a | n/a | n/a | n/a | n/a | n/a |
| **disorder** | n/a | 369 | 372 | n/a | n/a | n/a | n/a | n/a | n/a |
| **disorder** | n/a | 384 | 385 | n/a | n/a | n/a | n/a | n/a | n/a |

**Table S2.** This table includes pfam predicted domain of the *A. nidulans* protein Q5B1W7 (Uniprot identifier). A conserved lytic polysaccharide monooxygenase, cellulose-degrading domain (LPMO_10) as well as a carbohydrate/starch binding domain (CBM_20) with pfam accessions PF03067 and PF00686 are respectively predicted within Q5B1W7.

Table S3 - *Aspergillus nidulans* strains used in this study.

| **Strain ID** | **Modified site/s**[1] | **Genotype** | **Reference** |
|---|---|---|---|
| NID174/FGSC A4 | - | - | NCBI:txid227321 |
| NID2531 | Host/reference | *argB2*, *veA1*, *nkuA*Δ | In house collection |
| NID2134 | IS1[2] | *argB2; pyrG89; veA1; nkuA*Δ*; IS1::*P*gpdA-AN10118-RFP*-T*trpC::argB; IS4::*P*gpdA-mpaA-mCitrine*-T*trpC::AFpyrG* | In house collection |
| NID2838 | AN10909 | *argB2*, *veA1*, *nkuA*Δ, AN10909Δ | This study |
| NID2714 | AN2165 | *argB2*, *veA1*, *nkuA*Δ, AN2165Δ | This study |
| NID2721 | AN1566 | *argB2*, *veA1*, *nkuA*Δ, AN1566Δ | This study |
| NID2715 | AN0761 | *argB2*, *veA1*, *nkuA*Δ, AN0761Δ | This study |
| NID2733 | AN0134 | *argB2*, *veA1*, *nkuA*Δ, AN0134Δ | This study |
| NID2709 | AN2405 | *argB2*, *veA1*, *nkuA*Δ, AN2405Δ | This study |
| NID2713 | AN4663 | *argB2*, *veA1*, *nkuA*Δ, AN4663Δ | This study |
| NID2773 | AN4663 | *argB2*, *veA1*, *nkuA*Δ, AN4663Δ | This study |
| NID2774 | AN4663 | *argB2*, *veA1*, *nkuA*Δ, AN4663Δ | This study |
| NID2775 | AN4663 | *argB2*, *veA1*, *nkuA*Δ, AN4663Δ | This study |
| NID2723 | AN7375 | *argB2*, *veA1*, *nkuA*Δ, AN7375Δ | This study |
| NID2712 | AN10974 | *argB2*, *veA1*, *nkuA*Δ, AN10974Δ | This study |
| NID2745 | AN8945 | *argB2*, *veA1*, *nkuA*Δ, AN8945Δ | This study |
| NID2746 | AN9193 | *argB2*, *veA1*, *nkuA*Δ, AN9193Δ | This study |
| NID2747 | AN4625 | *argB2*, *veA1*, *nkuA*Δ, AN4625Δ | This study |
| NID2749 | AN5874 | *argB2*, *veA1*, *nkuA*Δ, AN5874Δ | This study |
| NID2750 | AN10700 | *argB2*, *veA1*, *nkuA*Δ, AN10700Δ | This study |
| NID2755 | AN5630 | *argB2*, *veA1*, *nkuA*Δ, AN5630Δ | This study |
| NID2757 | AN2406 | *argB2*, *veA1*, *nkuA*Δ, AN2406Δ | This study |
| NID2759 | AN9098 | *argB2*, *veA1*, *nkuA*Δ, AN9098Δ | This study |
| NID2802 | AN6094 | *argB2*, *veA1*, *nkuA*Δ, AN6094Δ | This study |
| NID2801 | AN3096 | *argB2*, *veA1*, *nkuA*Δ, AN3096Δ | This study |

| | | | |
|---|---|---|---|
| NID2761 | AN8849 | *argB2*, *veA1*, *nkuA*Δ, AN8849Δ | This study |
| NID2710 | AN3464 | *argB2*, *veA1*, *nkuA*Δ, AN3464Δ | This study |
| NID2825 | AN6713 | *argB2*, *veA1*, *nkuA*Δ, AN6713Δ | This study |
| NID2756 | AN4663 & IS5 | *argB2*, *veA1*, *nkuA*Δ, AN4663Δ, IS5::Ptef-AN4663-Ttef:: | This study |
| NID2751 | AN4663 & IS5 | *argB2*, *veA1*, *nkuA*Δ, IS5::Ptef-AN4663-Ttef:: | This study |
| NID2787 | AN4663 | *argB2*, *veA1*, *nkuA*Δ, AN4663::E340A:: | This study |
| NID2815 | AN4663 | *argB2*, *veA1*, *nkuA*Δ, AN4663::E340A:: | This study |
| NID2789 | AN4663 & IS5 | *argB2*, *veA1*, *nkuA*Δ, AN4663Δ, IS5::Ptef-*mRFP*-AN4663-Ttef:: | This study |
| NID2794 | AN4663 & IS5 | *argB2*, *veA1*, *nkuA*Δ, AN4663Δ, IS5::Ptef-AN4663-*mRFP*-Ttef:: | This study |
| NID2858 | AN4663 & IS5 | *argB2*, *veA1*, *nkuA*Δ, AN4663Δ, IS5::Pan4663-AN4663-Tan4663:: | This study |
| NID2851 | AN4663 & IS5 | argB2, veA1, nkuAΔ, AN4663Δ, IS5::Pan4663-*mRFP*-AN4663-Tan4663:: | This study |
| NID2852 | AN4663 & IS5 | *argB2*, *veA1*, *nkuA*Δ, AN4663Δ, IS5::Pan4663-*mRFP*-AN4663-Tan4663:: | This study |
| NID2853 | AN4663 & IS5 | *argB2*, *veA1*, *nkuA*Δ, AN4663Δ, IS5::Pan4663-*mRFP*-AN4663-Tan4663:: | This study |
| NID2843 | AN4663 & IS5 | *argB2*, *veA1*, *nkuA*Δ, AN4663Δ, IS5::Pan4663-Truncated_AN4663-Tan4663:: | This study |
| NID2844 | AN4663 & IS5 | *argB2*, *veA1*, *nkuA*Δ, AN4663Δ, IS5::Pan4663-Truncated_AN4663-Tan4663:: | This study |
| NID2845 | AN4663 & IS5 | *argB2*, *veA1*, *nkuA*Δ, AN4663Δ, IS5::Pan4663-Truncated_AN4663-Tan4663:: | This study |
| NID2857 | IS5 | *argB2*, *veA1*, *nkuA*Δ, IS5::Ptef-*mRFP*-Ttef:: | This Study |

[1] Genomic loci altered compared with NID2531

[2] IS1 for integration site 1 from Hansen et al, 2011 (Appl. Environ. Microbiol., 77 [2011]) ; IS5 is from from Holm et al, 2013 (PhD Thesis: Development and Implementation of Novel Genetic Tools for Investigation of Fungal Secondary Metabolism. Technical University of Denmark.), and Schalén et al., 2016 (Fungal Biol. Biotechnol. 3, 3).

Table S4 - *K. phaffii* strains used in this study.

| **Strain** | **Genotype** | **Source/Reference** |
|---|---|---|
| *E. coli* NEB5α | *fhuA2* Δ(*argF-lacZ*)U169 *phoA glnV44* Φ80 Δ(*lacZ*)M15 *gyrA96 recA1 relA1 endA1 thi-1 hsdR17* | a |
| *K. phaffii (P. pastoris)* GS115 | *his4* | b |
| *K. phaffii* GS115 α-MF$^{SP}$-LsAA9A Mut$^{+}$ | *aox1*::$P_{AOX1}$-α-MF$^{SP}$-LsAA9A-His6-*HIS4* | This study |
| *K. phaffii* GS115 Amy$^{SP}$-LsAA9A Mut$^{+}$ | *aox1*::$P_{AOX1}$-Amy$^{SP}$-LsAA9A-His6-*HIS4* | This study |
| *K. phaffii* GS115 Native$^{SP}$-LsAA9A Mut$^{+}$ | *aox1*::$P_{AOX1}$-Native$^{SP}$-LsAA9A-His6-*HIS4* | This study |
| *K. phaffii* GS115 α-MF$^{SP}$-LsAA9A Mut$^{+}$ | *aox1*::$P_{AOX1}$-α-MF$^{SP}$-LsAA9A-His6-*HIS4* + pBGP1-AN4663 | This study |
| *K. phaffii* GS115 Amy$^{SP}$-LsAA9A Mut$^{+}$ | *aox1*::$P_{AOX1}$-Amy$^{SP}$-LsAA9A-His6-*HIS4* + pBGP1-AN4663 | This study |
| *K. phaffii* GS115 Native$^{SP}$-LsAA9A Mut$^{+}$ | *aox1*::$P_{AOX1}$-Native$^{SP}$-LsAA9A-His6-*HIS4* + pBGP1-AN4663 | This study |
| *K. phaffii* GS115 Amy$^{SP}$-Tf10A Mut$^{+}$ | *aox1*::$P_{AOX1}$-Amy$^{SP}$-Tf10A-*HIS4* | This study |
| *K. phaffii* GS115 Amy$^{SP}$-Tf10A Mut$^{+}$ | *aox1*::$P_{AOX1}$-Amy$^{SP}$-Tf10A-*HIS4* + pBGP1-AN4663 | This study |

[a] NEB, Ipswich, MA, USA; [b]Thermo Fisher Scientific, Waltham, MA, USA

Table S5 - Plasmids used in this study

| **Plasmid** | **Description** | **Reference** |
|---|---|---|
| pLyGo-*Kp*-1 | Integrative vector into the AOX1 locus. Encodes the α-MF signal peptide for secretion. $Km^R$ (*E. coli*), *HIS4* (*K. phaffii*) | [1] |
| pLyGo-Kp-1-LsAA9A | Integrative vector into the AOX1 locus. Encodes the α-MF signal peptide for secretion and LsAA9A. $Km^R$ (*E. coli*), *HIS4* (*K. phaffii*) | [1] |
| pLyGo-*Kp*-2 | Integrative vector into the AOX1 locus. Encodes the Amy signal peptide for secretion. $Km^R$ (*E. coli*), *HIS4* (*K. phaffii*) | [1] |
| pLyGo-*Kp*-2-LsAA9A | Integrative vector into the AOX1 locus. Encodes the Amy signal peptide for secretion and LsAA9A. $Km^R$ (*E. coli*), *HIS4* (*K. phaffii*) | 1 |
| pLyGo-*Kp*-2-Tf10A | Integrative vector into the AOX1 locus. Encodes the Amy signal peptide for secretion and Tf10A. $Km^R$ (*E. coli*), *HIS4* (*K. phaffii*) | This study |
| pPIC9K-Native$^{SP}$-LsAA9A | Integrative vector into the AOX1 locus. Encodes the Native signal peptide for secretion of LsAA9A and LsAA9A. $Km^R$ (*E. coli*), *HIS4* (*K. phaffii*) | This study |
| pBGP1 | Episomal vector with a pGAP constitute promoter, $Amp^R$ (*E. coli*), $Zeocin^R$ (*K. phaffii*) | [2] |
| pFC331 | CRISPR-Cas9 vector, $Amp^R$ (*E. coli*), ORI, *argB* (*A. nidulans)*, AMA1. | [3] |
| pcAN10909 | CRISPR-Cas9 vector, $Amp^R$ (*E. coli*), ORI, *argB* (*A. nidulans)*, AMA1, sgRNA targeting AN10909 flanked by self-splicing tRNA. | This study |
| pcAN2165 | CRISPR-Cas9 vector, $Amp^R$ (*E. coli*), ORI, *argB* (*A. nidulans)*, AMA1, sgRNA targeting AN2165 flanked by self-splicing tRNA. | This study |
| pcAN1566 | CRISPR-Cas9 vector, $Amp^R$ (*E. coli*), ORI, *argB* (*A. nidulans)*, AMA1, sgRNA targeting AN1566 flanked by self-splicing tRNA. | This study |
| pcAN0761 | CRISPR-Cas9 vector, $Amp^R$ (*E. coli*), ORI, *argB* (*A. nidulans)*, AMA1, sgRNA targeting AN0761 flanked by self-splicing tRNA. | This study |
| pcAN0134 | CRISPR-Cas9 vector, $Amp^R$ (*E. coli*), ORI, *argB* (*A. nidulans)*, AMA1, sgRNA targeting AN0134 flanked by self-splicing tRNA. | This study |
| pcAN2405 | CRISPR-Cas9 vector, $Amp^R$ (*E. coli*), ORI, *argB* (*A. nidulans)*, AMA1, sgRNA targeting AN2405 flanked by self-splicing tRNA. | This study |

| | | |
|---|---|---|
| pcAN4663 | CRISPR-Cas9 vector, $Amp^R$ (*E. coli*), ORI, *argB* (*A. nidulans)*, AMA1, sgRNA targeting AN4663 flanked by self-splicing tRNA. | This study |
| pcAN7375 | CRISPR-Cas9 vector, $Amp^R$ (*E. coli*), ORI, *argB* (*A. nidulans)*, AMA1, sgRNA targeting AN7375 flanked by self-splicing tRNA. | This study |
| pcAN10974 | CRISPR-Cas9 vector, $Amp^R$ (*E. coli*), ORI, *argB* (*A. nidulans)*, AMA1, sgRNA targeting AN10974 flanked by self-splicing tRNA. | This study |
| pcAN8945 | CRISPR-Cas9 vector, $Amp^R$ (*E. coli*), ORI, *argB* (*A. nidulans)*, AMA1, sgRNA targeting AN8945 flanked by self-splicing tRNA. | This study |
| pcAN9193 | CRISPR-Cas9 vector, $Amp^R$ (*E. coli*), ORI, *argB* (*A. nidulans)*, AMA1, sgRNA targeting AN9193 flanked by self-splicing tRNA. | This study |
| pcAN3464 | CRISPR-Cas9 vector, $Amp^R$ (*E. coli*), ORI, *argB* (*A. nidulans)*, AMA1, sgRNA targeting AN3464 flanked by self-splicing tRNA. | This study |
| pcAN4625 | CRISPR-Cas9 vector, $Amp^R$ (*E. coli*), ORI, *argB* (*A. nidulans)*, AMA1, sgRNA targeting AN4625 flanked by self-splicing tRNA. | This study |
| pcAN6094 | CRISPR-Cas9 vector, $Amp^R$ (*E. coli*), ORI, *argB* (*A. nidulans)*, AMA1, sgRNA targeting AN6094 flanked by self-splicing tRNA. | This study |
| pcAN9098 | CRISPR-Cas9 vector, $Amp^R$ (*E. coli*), ORI, *argB* (*A. nidulans)*, AMA1, sgRNA targeting AN9098 flanked by self-splicing tRNA. | This study |
| pcAN3096 | CRISPR-Cas9 vector, $Amp^R$ (*E. coli*), ORI, *argB* (*A. nidulans)*, AMA1, sgRNA targeting AN3096 flanked by self-splicing tRNA. | This study |
| pcAN5874 | CRISPR-Cas9 vector, $Amp^R$ (*E. coli*), ORI, *argB* (*A. nidulans)*, AMA1, sgRNA targeting AN5874 flanked by self-splicing tRNA. | This study |
| pcAN6713 | CRISPR-Cas9 vector, $Amp^R$ (*E. coli*), ORI, *argB* (*A. nidulans)*, AMA1, sgRNA targeting AN6713 flanked by self-splicing tRNA. | This study |
| pcAN8849 | CRISPR-Cas9 vector, $Amp^R$ (*E. coli*), ORI, *argB* (*A. nidulans)*, AMA1, sgRNA targeting AN8849 flanked by self-splicing tRNA. | This study |
| pcAN10700 | CRISPR-Cas9 vector, $Amp^R$ (*E. coli*), ORI, *argB* (*A. nidulans)*, AMA1, sgRNA targeting AN10700 flanked by self-splicing tRNA. | This study |
| pcAN5630 | CRISPR-Cas9 vector, $Amp^R$ (*E. coli*), ORI, *argB* (*A. nidulans)*, AMA1, sgRNA targeting AN5630 flanked by self-splicing tRNA. | This study |
| pcAN2406 | CRISPR-Cas9 vector, $Amp^R$ (*E. coli*), ORI, *argB* (*A. nidulans)*, AMA1, sgRNA targeting AN2406 flanked by self-splicing tRNA. | This study |

| | | |
|---|---|---|
| pcIS5 | CRISPR-Cas9 vector, Amp$^R$ (*E. coli*), ORI, *argB* (*A. nidulans)*, AMA1, sgRNA targeting IS5 flanked by self-splicing tRNA. | in house collection |
| pcAN4663-PM-PS#2 | CRISPR-Cas9 vector, Amp$^R$ (*E. coli*), ORI, *argB* (*A. nidulans)*, AMA1, sgRNA targeting E340 of AN4663 flanked by self-splicing tRNA. | This study |
| pIS5-AN4663 | Amp$^R$ (*E. coli*), ORI. Integration vector for insertion of Ptef, AN4663, and Ttef in integration site 5 | This study |
| pIS5-AN4663-N-tag-mRFP | Amp$^R$ (*E. coli*), ORI. Integration vector for insertion of AN4663 with Ptef promoter and Ttef terminator and N-terminal tagged with mRFP in integration site 5 | This study |
| pIS5-AN4663-C-tag-mRFP | Amp$^R$ (*E. coli*), ORI. Integration vector for insertion of AN4663 with Ptef promoter and Ttef terminator and C-terminal tagged with mRFP in integration site 5 | This study |
| pIS5-Pan4663-AN4663-Tan4663 | Amp$^R$ (*E. coli*), ORI. Integration vector for insertion of AN4663 with native promoter and terminator in integration site 5 | This study |
| pIS5-AN4663-TruncV2 Native | Amp$^R$ (*E. coli*), ORI. Integration vector for insertion of truncated AN4663 after membrane region with native promoter and terminator in integration site 5 | This study |
| pIS5-Ptef-mRFP-Ttef | Amp$^R$ (*E. coli*), ORI. Integration vector for insertion of Ptef, mRFP, and Ttef in integration site 5 | This study |

1. C. Hernández-Rollán, K. B. Falkenberg, M. Rennig, A. B. Bertelsen, J. Ø. Ipsen, S. Brander, D. O. Daley, K. S. Johansen, M. H. H. Nørholm, LyGo: A Platform for Rapid Screening of Lytic Polysaccharide Monooxygenase Production. ACS Synthetic Biology. 10, 897–906 (2021).

2. C. C. Lee, T. G. Williams, D. W. S. Wong, G. H. Robertson, An episomal expression vector for screening mutant gene libraries in *Pichia pastoris*. Plasmid. 54, 80–85 (2005).

3. C. S. Nødvig, J. B. Nielsen, M. E. Kogle, U. H. Mortensen, A CRISPR-Cas9 System for Genetic Engineering of Filamentous Fungi. PLOS ONE. 10, e0133085 (2015).

Table S6 – Oligonucleotides and target sequences used in this study.

Color code for: annealing sequence, protospacer sequence, thymine → uracil substitution, overhangs for PacI/Nt.BbvCI cassette

| ID | Purpose | Sequence (5' à 3') |
|---|---|---|
| **Primers CRISPR/Cas9-vector construction** | | |
| CSN438-Afum-U3p-fwd | Amplifying CRISPR fragment with promoter | GGGTTTAAUGATCACATAGATGCTCGGTTGACA |
| CSN790-U3-term-rv | Amplifying CRISPR fragment with terminator | GGTCTTAAUACCCTGAGAAGATAGATGTGAATGTG |
| tRNA-gRNA328-rv | Amplifying CRISPR fragment with PS tails targeting AN10909 | AGGACATTAUGCACTCGTGCATCATCCGTGAATCGAAC |
| gRNA328-fwd | Amplifying CRISPR fragment with PS tails targeting AN10909 | ATAATGTCCUTCAGTTTTAGAGCTAGAAATAGCAAGTTAAA |
| tRNA-gRNA329-rv | Amplifying CRISPR fragment with PS tails targeting AN10909 | ATGCAGGTGGUGGGTGCATCATCCGTGAATCGAAC |
| gRNA329-fwd | Amplifying CRISPR fragment with PS tails targeting AN10909 | ACCACCTGCAUTAGTTCGTTTTAGAGCTAGAAATAGCAAGTTAAA |
| tRNA-gRNA330-rv | Amplifying CRISPR fragment with PS tails targeting AN2165 | ATCTGCTCATGGAAAUTTTGCATCATCCGTGAATCGAAC |
| gRNA330-fwd | Amplifying CRISPR fragment with PS tails targeting AN2165 | ATTTCCATGAGCAGAUTCGTTTTAGAGCTAGAAATAGCAAGTTAAA |
| tRNA-gRNA331-rv | Amplifying CRISPR fragment with PS tails targeting AN2165 | AGTTCATTGCTUCGCGTGCATCATCCGTGAATCGAAC |
| gRNA331-fwd | Amplifying CRISPR fragment with PS tails targeting AN2165 | AAGCAATGAACUGCTCGTTTTAGAGCTAGAAATAGCAAGTTAAA |
| tRNA-gRNA332-rv | Amplifying CRISPR fragment with PS tails targeting AN1566 | ACCAGTTGTCCUTATGCATCATCCGTGAATCGAAC |
| gRNA332-fwd | Amplifying CRISPR fragment with PS tails targeting AN1566 | AGGACAACTGGUCATCCGGTTTTAGAGCTAGAAATAGCAAGTTAAA |
| tRNA-gRNA333-rv | Amplifying CRISPR fragment with PS tails targeting AN1566 | AACATACCCUAGCCTGCATCATCCGTGAATCGAAC |
| gRNA333-fwd | Amplifying CRISPR fragment with PS tails targeting AN1566 | AGGGTATGTUAGGCTTGTTTTAGAGCTAGAAATAGCAAGTTAAA |
| tRNA-gRNA338-rv | Amplifying CRISPR fragment with PS tails targeting AN0761 | ATCTCAGCACCUCCCGATTGCATCATCCGTGAATCGAAC |
| gRNA338-fwd | Amplifying CRISPR fragment with PS tails targeting AN0761 | AGGTGCTGAGAUGAGTTTTAGAGCTAGAAATAGCAAGTTAAA |
| tRNA-gRNA339-rv | Amplifying CRISPR fragment with PS tails targeting AN0761 | AGGGCAGAAGGGUGATGCATCATCCGTGAATCGAAC |
| gRNA339-fwd | Amplifying CRISPR fragment with PS tails targeting AN0761 | ACCCTTCTGCCCUATTTGGTTTTAGAGCTAGAAATAGCAAGTTAAA |
| tRNA-gRNA340-rv | Amplifying CRISPR fragment with PS tails targeting AN0134 | AGATTCCAUACTGTGCATCATCCGTGAATCGAAC |
| gRNA340-fwd | Amplifying CRISPR fragment with PS tails targeting AN0134 | ATGGAATCUGAAAGCGGTTTTAGAGCTAGAAATAGCAAGTTAAA |
| tRNA-gRNA341-rv | Amplifying CRISPR fragment with PS tails targeting AN0134 | AATGCTTGAGUGTATGCATCATCCGTGAATCGAAC |
| gRNA341-fwd | Amplifying CRISPR fragment with PS tails targeting AN0134 | ACTCAAGCATUAAGGATGTTTTAGAGCTAGAAATAGCAAGTTAAA |
| tRNA-gRNA-AN4663-rv(B) | Amplifying CRISPR fragment with PS tails targeting AN4663 | ACCAGCTTCTTGGUAGCTGCATCATCCGTGAATCGAAC |
| gRNA_AN4663-fwd(B) | Amplifying CRISPR fragment with PS tails targeting AN4663 | ACCAAGAAGCTGGUAGTGTTTTAGAGCTAGAAATAGCAAGTTAAA |
| tRNA-gRNA-AN4663-rv(E) | Amplifying CRISPR fragment with PS tails targeting AN4663 | ACAGTCCTCAUAATCTGCATCATCCGTGAATCGAAC |
| gRNA_AN4663-fwd(E) | Amplifying CRISPR fragment with PS tails targeting AN4663 | ATGAGGACTGUTTTGCGTTTTAGAGCTAGAAATAGCAAGTTAAA |
| tRNA-gRNA-AN3464-rv(B) | Amplifying CRISPR fragment with PS tails targeting AN3464 | AGTACCACGCCAUCGTTGCATCATCCGTGAATCGAAC |
| gRNA_AN3464-fwd(B) | Amplifying CRISPR fragment with PS tails targeting AN3464 | ATGGCGTGGTACUGTTCGTTTTAGAGCTAGAAATAGCAAGTTAAA |
| tRNA-gRNA-AN3464-rv(E) | Amplifying CRISPR fragment with PS tails targeting AN3464 | AATGCGGTACUGATGCATCATCCGTGAATCGAAC |
| gRNA_AN3464-fwd(E) | Amplifying CRISPR fragment with PS tails targeting AN3464 | AGTACCGCATUCGGCGGTGTTTTAGAGCTAGAAATAGCAAGTTAAA |
| tRNA-gRNA-AN2405-rv(B) | Amplifying CRISPR fragment with PS tails targeting AN2405 | ATTGCTTTTGCUCTGCATCATCCGTGAATCGAAC |

| | | |
|---|---|---|
| gRNA_AN2405-fwd(B) | Amplifying CRISPR fragment with PS tails targeting AN2405 | AGCAAAAGCAAUCCACCTTGTTTTAGAGCTAGAAATAGCAAGTTAAA |
| tRNA-gRNA-AN2405-rv(E) | Amplifying CRISPR fragment with PS tails targeting AN2405 | ACCTTCTTTGUCAACGTCTGCATCATCCGTGAATCGAAC |
| gRNA_AN2405-fwd(E) | Amplifying CRISPR fragment with PS tails targeting AN2405 | ACAAAGAAGGUCGGTTTTAGAGCTAGAAATAGCAAGTTAAA |
| tRNA-gRNA-AN7375-rv(B) | Amplifying CRISPR fragment with PS tails targeting AN7375 | AGCTCGACTGAUTTCTTTGCATCATCCGTGAATCGAAC |
| gRNA_AN7375-fwd(B) | Amplifying CRISPR fragment with PS tails targeting AN7375 | ATCAGTCGAGCUCTCGTTTTAGAGCTAGAAATAGCAAGTTAAA |
| tRNA-gRNA-AN7375-rv(E) | Amplifying CRISPR fragment with PS tails targeting AN7375 | AGGCAACACCGUTTGCATCATCCGTGAATCGAAC |
| gRNA_AN7375-fwd(E) | Amplifying CRISPR fragment with PS tails targeting AN7375 | ACGGTGTTGCCUCTCAGTTGTTTTAGAGCTAGAAATAGCAAGTTAAA |
| tRNA-gRNA-AN10974-rv(B) | Amplifying CRISPR fragment with PS tails targeting AN10974 | ATGTCGACTATGUCAATGCATCATCCGTGAATCGAAC |
| gRNA_AN10974-fwd(B) | Amplifying CRISPR fragment with PS tails targeting AN10974 | ACATAGTCGACAUTCGCGTTTTAGAGCTAGAAATAGCAAGTTAAA |
| tRNA-gRNA-AN10974-rv(E) | Amplifying CRISPR fragment with PS tails targeting AN10974 | ACCGCTTCCAAACUCATGCATCATCCGTGAATCGAAC |
| gRNA_AN10974-fwd(E) | Amplifying CRISPR fragment with PS tails targeting AN10974 | AGTTTGGAAGCGGUTCCGGTTTTAGAGCTAGAAATAGCAAGTTAAA |
| tRNA-gRNA-AN9193-rv1 | Amplifying CRISPR fragment with PS tails targeting AN9193 | ATGGCTGACGCUGCATGCATCATCCGTGAATCGAAC |
| gRNA-AN9193-fwd1 | Amplifying CRISPR fragment with PS tails targeting AN9193 | AGCGTCAGCCAUGGTCAGTTTTAGAGCTAGAAATAGCAAGTTAAA |
| tRNA-gRNA-AN9193-rv2 | Amplifying CRISPR fragment with PS tails targeting AN9193 | ACAACATTGAACCUGCTGCATCATCCGTGAATCGAAC |
| gRNA-AN9193-fwd2 | Amplifying CRISPR fragment with PS tails targeting AN9193 | AGGTTCAATGTTGUGGTCGTTTTAGAGCTAGAAATAGCAAGTTAAA |
| tRNA-gRNA-AN8945-rv1 | Amplifying CRISPR fragment with PS tails targeting AN8945 | ACCGACAAAAUATGCATCATCCGTGAATCGAAC |
| gRNA-AN8945-fwd1 | Amplifying CRISPR fragment with PS tails targeting AN8945 | ATTTTGTCGGUCGCAATGGGTTTTAGAGCTAGAAATAGCAAGTTAAA |
| tRNA-gRNA-AN8945-rv2 | Amplifying CRISPR fragment with PS tails targeting AN8945 | ACTCCGGCTTCUGCCTGCATCATCCGTGAATCGAAC |
| gRNA-AN8945-fwd2 | Amplifying CRISPR fragment with PS tails targeting AN8945 | AGAAGCCGGAGUAGAGCGTTTTAGAGCTAGAAATAGCAAGTTAAA |
| tRNA-gRNA-AN4625-rv1 | Amplifying CRISPR fragment with PS tails targeting AN4625 | AGTAGGTGTCTUGAATCTGCATCATCCGTGAATCGAAC |
| gRNA-AN4625-fwd1 | Amplifying CRISPR fragment with PS tails targeting AN4625 | AAGACACCTACUCCGGTTTTAGAGCTAGAAATAGCAAGTTAAA |
| tRNA-gRNA-AN4625-rv2 | Amplifying CRISPR fragment with PS tails targeting AN4625 | ACAGCATAGAUCTGTGCATCATCCGTGAATCGAAC |
| gRNA-AN4625-fwd2 | Amplifying CRISPR fragment with PS tails targeting AN4625 | ATCTATGCTGUCAAGGGGTTTTAGAGCTAGAAATAGCAAGTTAAA |
| tRNA-gRNA-AN6094-rv1 | Amplifying CRISPR fragment with PS tails targeting AN6094 | ATGGGTTGTUGTGAAGGTGCATCATCCGTGAATCGAAC |
| gRNA-AN6094-fwd1 | Amplifying CRISPR fragment with PS tails targeting AN6094 | AACAACCCAUAAAGTTTTAGAGCTAGAAATAGCAAGTTAAA |
| tRNA-gRNA-AN6094-rv2 | Amplifying CRISPR fragment with PS tails targeting AN6094 | AAAATCACGUACAACCAGTGCATCATCCGTGAATCGAAC |
| gRNA-AN6094-fwd2 | Amplifying CRISPR fragment with PS tails targeting AN6094 | ACGTGATTTUCCGTTTTAGAGCTAGAAATAGCAAGTTAAA |
| tRNA-gRNA-AN9098-rv1 | Amplifying CRISPR fragment with PS tails targeting AN9098 | AGCTGAGUAAATTGGATTGCATCATCCGTGAATCGAAC |
| gRNA-AN9098-fwd1 | Amplifying CRISPR fragment with PS tails targeting AN9098 | ACTCAGCUTCAGTTTTAGAGCTAGAAATAGCAAGTTAAA |
| tRNA-gRNA-AN9098-rv2 | Amplifying CRISPR fragment with PS tails targeting AN9098 | ATCCCGGTTAUATCCATATCTGCATCATCCGTGAATCGAAC |
| gRNA-AN9098-fwd2 | Amplifying CRISPR fragment with PS tails targeting AN9098 | ATAACCGGGAUGTTTTAGAGCTAGAAATAGCAAGTTAAA |
| tRNA-gRNA-AN3096-rv1 | Amplifying CRISPR fragment with PS tails targeting AN3096 | AGTAGTCGCCAUGTCCGCTGCATCATCCGTGAATCGAAC |
| gRNA-AN3096-fwd1 | Amplifying CRISPR fragment with PS tails targeting AN3096 | ATGGCGACTACUGCGTTTTAGAGCTAGAAATAGCAAGTTAAA |
| tRNA-gRNA-AN3096-rv2 | Amplifying CRISPR fragment with PS tails targeting AN3096 | AGTTCTGGCTUCCCCTTTCGTGCATCATCCGTGAATCGAAC |

| | | |
|---|---|---|
| gRNA-AN3096-fwd2 | Amplifying CRISPR fragment with PS tails targeting AN3096 | AAGCCAGAACUGTTTTAGAGCTAGAAATAGCAAGTTAAA |
| tRNA-gRNA-AN2406-rv1 | Amplifying CRISPR fragment with PS tails targeting AN2406 | ATTGATGTTTTCUGCCCTTGCATCATCCGTGAATCGAAC |
| gRNA-AN2406-fwd1 | Amplifying CRISPR fragment with PS tails targeting AN2406 | AGAAAACATCAAUATGTTTTAGAGCTAGAAATAGCAAGTTAAA |
| tRNA-gRNA-AN2406-rv2 | Amplifying CRISPR fragment with PS tails targeting AN2406 | ACCTTCGCACCUTGTATATGCATCATCCGTGAATCGAAC |
| gRNA-AN2406-fwd2 | Amplifying CRISPR fragment with PS tails targeting AN2406 | AGGTGCGAAGGUAGGTTTTAGAGCTAGAAATAGCAAGTTAAA |
| tRNA-gRNA-AN5630-rv1 | Amplifying CRISPR fragment with PS tails targeting AN5630 | ATTTTCGGTGACCAUGTGCATCATCCGTGAATCGAAC |
| gRNA-AN5630-fwd1 | Amplifying CRISPR fragment with PS tails targeting AN5630 | ATGGTCACCGAAAAUGGCTGTTTTAGAGCTAGAAATAGCAAGTTAAA |
| tRNA-gRNAAN5630-rv2 | Amplifying CRISPR fragment with PS tails targeting AN5630 | AGTGAGTAAATTGUGGTGCATCATCCGTGAATCGAAC |
| gRNA-AN5630-fwd2 | Amplifying CRISPR fragment with PS tails targeting AN5630 | ACAATTTACTCACUCTTGGTTTTAGAGCTAGAAATAGCAAGTTAAA |
| tRNA-gRNA-AN5874-rv1 | Amplifying CRISPR fragment with PS tails targeting AN5874 | ATCTGGTCAGGUTGTGCATCATCCGTGAATCGAAC |
| gRNA-AN5874-fwd1 | Amplifying CRISPR fragment with PS tails targeting AN5874 | ACCTGACCAGAUATTTGTGTTTTAGAGCTAGAAATAGCAAGTTAAA |
| tRNA-gRNA-AN5874-rv2 | Amplifying CRISPR fragment with PS tails targeting AN5874 | ATCTCGGCTTGUCGGGTCTGCATCATCCGTGAATCGAAC |
| gRNA-AN5874-fwd2 | Amplifying CRISPR fragment with PS tails targeting AN5874 | ACAAGCCGAGAUATGTTTTAGAGCTAGAAATAGCAAGTTAAA |
| tRNA-gRNA-AN6713-rv1 | Amplifying CRISPR fragment with PS tails targeting AN6713 | ACGCGATTTCUACAAATGCATCATCCGTGAATCGAAC |
| gRNA-AN6713-fwd1 | Amplifying CRISPR fragment with PS tails targeting AN6713 | AGAAATCGCGUGCCAGTTTTAGAGCTAGAAATAGCAAGTTAAA |
| tRNA-gRNA-AN6713-rv2 | Amplifying CRISPR fragment with PS tails targeting AN6713 | AGTTTCCTCTUCATGCATCATCCGTGAATCGAAC |
| gRNA-AN6713-fwd2 | Amplifying CRISPR fragment with PS tails targeting AN6713 | AAGAGGAAACUGGAGGGGGTTTTAGAGCTAGAAATAGCAAGTTAAA |
| tRNA-gRNA-AN8849-rv1 | Amplifying CRISPR fragment with PS tails targeting AN8849 | AGTTGCTCTCUTTGCATCATCCGTGAATCGAAC |
| gRNA-AN8849-fwd1 | Amplifying CRISPR fragment with PS tails targeting AN8849 | AGAGAGCAACUAGCACGCAGTTTTAGAGCTAGAAATAGCAAGTTAAA |
| tRNA-gRNA-AN8849-rv2 | Amplifying CRISPR fragment with PS tails targeting AN8849 | ATTAATCACTATCUGATTGCATCATCCGTGAATCGAAC |
| gRNA-AN8849-fwd2 | Amplifying CRISPR fragment with PS tails targeting AN8849 | AGATAGTGATTAAUTTAGTTTTAGAGCTAGAAATAGCAAGTTAAA |
| tRNA-gRNA-AN10700-rv1 | Amplifying CRISPR fragment with PS tails targeting AN10700 | AGTTTCTCGGUGGATGAATGCATCATCCGTGAATCGAAC |
| gRNA-AN10700-fwd1 | Amplifying CRISPR fragment with PS tails targeting AN10700 | ACCGAGAAACUAAGTTTTAGAGCTAGAAATAGCAAGTTAAA |
| tRNA-gRNA-AN10700-rv2 | Amplifying CRISPR fragment with PS tails targeting AN10700 | ACTTCTTGAAAGCCUCTGCATCATCCGTGAATCGAAC |
| gRNA-AN10700-fwd2 | Amplifying CRISPR fragment with PS tails targeting AN10700 | AGGCTTTCAAGAAGUTTAGGTTTTAGAGCTAGAAATAGCAAGTTAAA |
| tRNA-gRNA-AN4663-PM-rv-2 | Amplifying CRISPR fragment with PS tails targeting AN4663 E340 | ATTTCGACGAUGGTCTGCATCATCCGTGAATCGAAC |
| gRNA-AN4663-PM-fwd-2 | Amplifying CRISPR fragment with PS tails targeting AN4663 E340 | ATCGTCGAAAUCGACCGTTTTAGAGCTAGAAATAGCAAGTTAAA |
| **Primers for Integration-vector construction** | | |
| IS5 P1 RV | Backbone to IS5 tails | ACATGGCAGUCGGCCGCATTTAAATCCCC |
| IS5 P1 FW | Backbone to IS5 tails | ATCTGGACTUAGCGCGGCCGCAAATTTAAAT |
| IS5 pAC1 DS RV | IS5 DS with Backbone tail | AAGTCCAGAUTTACTCAATTCAGGTCTG |
| IS5 pAC1 US FW | IS5 US with Backbone tail | ACTGCCATGUGTCCAAGCC |
| IS5 pAC1 US RV | IS5 US with PacI tail | AGGGTTUAATTAAGACCTCAGCTCGCTGAGTGTGAGTCTGACT |
| IS5 pAC1 DS FW | IS5 DS with PacI tail | AAACCCUCAGCGCATGGCAATCAAGTCCCTGT |
| Ptef-FU-PacI Up | Amplifying Ptef fragment with PacI tail | GGGTTTAAUCGAGACAGCAGAATCACCG |
| Ptef-RU-start two | Amplifying Ptef fragment | ATGGTGAAGGUTGTGTTATGTTTTG |
| Ttef-FU-end A | Amplifying Ttef fragment | AGCGGACAUTCGATTTATGC |
| Ttef-RU-PacI Dw | Ampligying Ttef fragment with PacI tail | GGTCTTAAUGTATTGGGATGAATTTTGTATGC |
| P1 FW | General Backbone to IS tail | ACGCATCGGUAGCGCGGCCGCAAATTTAAAT |

| | | |
|---|---|---|
| P1 RV | General Backbone to IS tail | AGGTCGTCCUCGGCCGCATTTAAATCCCC |
| AN4663-FU-Ptef | Amplifying AN4663 with Ptef tail | ACCTTCACCAUGGCGCCATTCAGGAGC |
| AN4663-RV-Ttef | Amplifying AN4663 with Ttef tail | ATGTCCGCUACCACCCCTCCCAGACC |
| AN4663-RV-mRFP v2 | Amplifying AN4553 with C terminal mRFP tail | AGGCCAUCCACCCCTCCCAGACCCTAT |
| AN4663-FW-mRFP | Amplifying AN4553 with N terminal mRFP tail | ATGGCGCCAUTCAGGAGCAT |
| mRFP-RV-AN4663 | Amplifying mRFP as C terminal with AN4663 tail | ATGGCGCCAUGGCGCCGGTGGAGTGG |
| mRFP-FW-AN4663 | Amplifying mRFP as N terminal with AN4663 tail | ATGGCCUCCTCCGAGGAC |
| AN4663-truncv2-FU | Amplifying Truncated AN4663 without transmembrane region | ACCTTCACCAUGGCCCTCAACAACATCATCGC |
| ISan4663-P1-DS-RU | Amplifying ISan4663 DS with Backbone tail | ACCGATGCGUGAGGTCGAGACATTAGATATGCAAGG |
| ISan4663-P1-US-FW | Amplifying ISan4663 US with Backbone tail | AGGACGACCUCTTCAACGCACCATTCAGGCT |
| ISan4663-PacI-US-RU | Amplifying ISan4663 US with PacI tail | AGGGTTUAATTAAGACCTCAGCGTTACTATTTGGTGATCAGGCGCA |
| ISan4663-PacI-DS-FW | Amplifying ISan4663 DS with PacI tail | AAACCCUCAGCATGTGTTTTTCCTGACCCAGGG |
| **Primers for DNA insertions & gene deletion check** | | |
| ANIS5-ChkUp-F2 | Checking for insertion in IS5 | CGTACTCTGTCCCACCAACC |
| ANIS5-ChkDw-R2 | Checking for insertion in IS5 | CCGCGTTATCCCTTCCTGC |
| AN-S53-Chk-Gap-F | Checking for insertion in IS5 | CTCCTTCCTAGCCTTGCAACTCC |
| AN-S53-Chk-Gap-R | Checking for insertion in IS5 | CCTAGCTCATTCCTCAGTCCGTC |
| AN-S53-Chk-Up-F | Checking for insertion in IS5 | GATTGCATGGTTGGATCTGGATG |
| AN-S53-Chk-Dw-R | Checking for insertion in IS5 | GGATGGCGAATCGTGAGCG |
| AN10909-Chk-FW | Checking for deletion of AN10909 | GGCTTTCGCCGTGTTGTGA |
| AN10909-Chk-RV | Checking for deletion of AN10909 | GGTGGTTGCAGTCCGTGA |
| AN2165-Chk-FW | Checking for deletion of AN2165 | CGCAGCGGTCAAATATCCC |
| AN2165-Chk-RV | Checking for deletion of AN2165 | GGGCGTTGGGTATTCTTTGC |
| AN1566-Chk-FW | Checking for deletion of AN1566 | GGACCTATACCTAGTCCCCC |
| AN1566-Chk-RV | Checking for deletion of AN1566 | AAATTGAGCGCCGCAAGC |
| AN0761-Chk-FW | Checking for deletion of AN0761 | ACGGTCCAGGACAAACGAC |
| AN0761-Chk-RV | Checking for deletion of AN0761 | CTTCTCTGGCTCTCACTGGG |
| AN0134-Chk-FW | Checking for deletion of AN0134 | CGACGATTTCGCTCGAAGTG |
| AN0134-Chk-RV | Checking for deletion of AN0134 | TAGTCCAACCCAGCTGCAT |
| AN4663-Chk-F | Checking for deletion of AN4663 | TTGCGGGGTGAAGATGGTAG |
| AN4663-Chk-R | Checking for deletion of AN4663 | ATGGGAATGGTGGGGTTGTT |
| AN3464-Chk-F | Checking for deletion of AN3464 | GCGTGATGGCGTAGGATGTA |
| AN3464-Chk-R | Checking for deletion of AN3464 | CCGTTTGGTGGAGGTCATCA |
| AN2405-Chk-F | Checking for deletion of AN2405 | GGAGGTGGTCTCGTTTGGAG |
| AN2405-Chk-R | Checking for deletion of AN2405 | TCAGCAGCAATTACCCGTCA |
| AN7375-Chk-F | Checking for deletion of AN7375 | ATTTCGGTTATGCGGGGAGC |
| AN7375-Chk-R | Checking for deletion of AN7375 | ACAATCACTTCGCCGTTCCA |
| AN10974-Chk-F | Checking for deletion of AN10974 | GGGCTGCTCGTATCTTACCC |
| AN10974-Chk-R | Checking for deletion of AN10974 | CCAATTGAAGTCCGTCTGCG |
| AN9193-Chk-F | Checking for deletion of AN9193 | GCAGAGTCTCCGCTTATGGT |
| AN9193-Chk-R | Checking for deletion of AN9193 | TCATATGCGACACGCAGGATT |
| AN8945-Chk-F | Checking for deletion of AN8945 | AGACGGAAGTGCCGGAAAC |
| AN8945-Chk-R | Checking for deletion of AN8945 | CTGGAAGCCGTCCAAGGTT |
| AN4625-Chk-F | Checking for deletion of AN4625 | GAGAATAAGCCGTGCGCAGC |
| AN4625-Chk-R | Checking for deletion of AN4625 | AGCCCTAGGCTGGAAGTAAG |
| AN6094-Chk-F | Checking for deletion of AN6094 | AGGCGTTTCAAGGTGTTACTTCG |
| AN6094-Chk-R | Checking for deletion of AN6094 | CCAACTCGGTAACATGATCTCCTC |
| AN9098-Chk-F | Checking for deletion of AN9098 | CTCTTCCTCGTTCTTTTCGCGC |
| AN9098-Chk-R | Checking for deletion of AN9098 | TTCTGCCCGTCGCGGC |
| AN3096-Chk-F | Checking for deletion of AN3096 | CTTATCAATAGCACAGCTGAAAGAAAGGG |
| AN3096-Chk-R | Checking for deletion of AN3096 | CAGTTGAGGAACAACGAAAGAAATGG |
| AN2406-Chk-F | Checking for deletion of AN2406 | TCTATTGAAGAGGTGGGAGCTGG |
| AN2406-Chk-R | Checking for deletion of AN2406 | GGTCGCGGAATGCTGGATAT |
| AN5630-Chk-F | Checking for deletion of AN5630 | CAGTCTCCGTACTTCATGGCT |
| AN5630-Chk-R | Checking for deletion of AN5630 | GATGAGGGGCTTGTGGAGAT |
| AN5874-Chk-F | Checking for deletion of AN5874 | ATTTGCGCAGGGAAAGGCTAC |
| AN5874-Chk-R | Checking for deletion of AN5874 | CAGCGCTCAACAGAGATGTTG |
| AN6713-Chk-F | Checking for deletion of AN6713 | ACCTTTCATGTGCTCTTCGAGG |
| AN6713-Chk-R | Checking for deletion of AN6713 | AGTCGTCCCTCTAATCACGC |
| AN8849-Chk-F | Checking for deletion of AN8849 | ATTTTGCGACTGCTTCCCCG |

| | | |
|---|---|---|
| AN8849-Chk-R | Checking for deletion of AN8849 | ATGAGCTCCTCTCATTGCCTTC |
| AN10700-Chk-F | Checking for deletion of AN10700 | ACGATGAGAGCTCCATCAAAGG |
| AN10700-Chk-R | Checking for deletion of AN10700 | CTGAACGTCTTGGCCAGAAGAT |
| **Oligonucleotides for gene editing** | | |
| GE-Oligo-328-329 | Reparing dsb to delete the gene AN10909 | ACACCACCACGATTAGTTTGCGAGTGCATAGTCAATCCGGCTAGCGGGATCCATTAGAGC |
| GE-Oligo-330-331 | Reparing dsb to delete the gene AN2165 | CCCTCTTAGAAGCTTCCAACTGCTCGGAGCCTTTCGGACGGACGAATAATGAATGATGCA |
| GE-Oligo-332-333 | Reparing dsb to delete the gene AN1566 | CGCCTAGTAACAACCGGGGCCCCGTGAACCGAGACTGTATTAGTAGATGTTCTGCGCATA |
| GE-Oligo-338-339 | Reparing dsb to delete the gene AN0761 | CACATCTCGGTGATACGGATCGCTCAGGGAAACATGGACGACCGAGGGTGAGCGAAAAGC |
| GE-Oligo-340-341 | Reparing dsb to delete the gene AN0134 | GTCTTCTTCCGTTTTATATCAGGTGTCAGTCGGAGAAAGTCTGCTTTAACTCGTTCCGGA |
| GE-Oligo-AN4663 | Reparing dsb to delete the gene AN4663 | TCTGTCTGCGCCTGATCACCAAATAGTAACATGTGTTTTTCCTGACCCAGGGCGAAAGTG |
| GE-Oligo-AN3464 | Reparing dsb to delete the gene AN3464 | CTAAAACAAACGACCAGAAGATATACAACGAGACCTGAGTTCTGCGATATGCTATGGAAT |
| GE-Oligo-AN2405 | Reparing dsb to delete the gene AN2405 | TGTTTTGTATAGAAGACGGATACGCTCACATGCCGCTATATCCGCTGCTCGTCATAGTAT |
| GE-Oligo-AN7375 | Reparing dsb to delete the gene AN7375 | GGGATCCTAGAGGTCAAAAATCAATCGACGGCTGCTGTTGGTGAATCTGTTATCAGAAGT |
| GE-Oligo-AN10974 | Reparing dsb to delete the gene AN10974 | AAGCTTCCGTTCGTCTGCCAAATAATCAACTCATGTAACAAAACCATATTAACTCATGGG |
| GE-Oligo--AN9193 | Reparing dsb to delete the gene AN9193 | AACAAGCCTTCAAACCTCCTAATAAACTCCCTCGCCGTACCAACCACTAGTTATATTCGG |
| GE-Oligo--AN8945 | Reparing dsb to delete the gene AN8945 | TCTTTTGATTTTACTATTTTGTCGGTCGCAAGCTGGCTTATTGTGGCTTCAGTCATCGTA |
| GE-Oligo--AN4625 | Reparing dsb to delete the gene AN4625 | AGCCTTCATTCTAATCCCAATTTATCGAGAACAGAACTCTACCGTACGTGCGCCAGATTT |
| GE-Oligo--AN6094 | Reparing dsb to delete the gene AN6094 | CCATCATTTATCTACAAGCTTGTCTACGAATCTCGCGTCTGGTTGTACGTGATTTTCCGG |
| GE-Oligo--AN9098 | Reparing dsb to delete the gene AN9098 | AGCTACAGAGAATCCAATTTACTCAGCTTCACGAAGTGATATATGATTCGGGAAAAAGCG |
| GE-Oligo--AN3096 | Reparing dsb to delete the gene AN3096 | GACTGGGTTTTTTCCCCCGCAGTAGTCGCCATATTAAGCTTTAAAATAAAAGTTTTGCTT |
| GE-Oligo-AN2406 | AN2406 | ATCCCCTCGCACAGGGCAGAAAACATCAATAGGTGATTCGGAAGAGTTGTATATACGGAC |
| GE-Oligo-AN5630 | AN5630 | ATTTGTCAGTACCAGGCATGGTCACCGAAAGTAAATTGTGGCGGACTTGCAATCTGCTCT |
| GE-Oligo-AN5874 | Reparing dsb to delete the gene AN5874 | AGAAAATACTTCAAGCCTTGACAGGCGATAGCGCGATGAATGGTGAAAATATCACCAAAC |
| GE-Oligo-AN6713 | Reparing dsb to delete the gene AN6713 | GCAATCACTTGCTGATATAGCTATTGCACCACTACTAGGTAGCTTTTTTGTCCGTATCAC |
| GE-Oligo-AN8849 | Reparing dsb to delete the gene AN8849 | TCCACATACGGAAGAGAGCAACTAGCACGCTTTATGGAAATACATGTATGCATAACAGAT |
| GE-Oligo-AN10700 | Reparing dsb to delete the gene AN10700 | CTTCCCAAACGTTCATCCACCGAGAAACTAGTTGACGATATGTTTTGATTTGATACATAA |
| GE-oligo AN4663 PM E340A | Reparing dsb and mutate AN4663 E340A | GAATCGACACGACCATCGTCGCCATCGACCCGGTTGTCCACAAGT |
| **Primers for sequencing** | | |
| pFC902sgRNA-Seq-F | To sequence the tRNA, protospacer, gRNA and tRNA | CATCGCATCCTCATGAAGTC |
| AN4663-PM-Seq-Fw | To sequence for checking the point mutation AN4663 E340A | GTCATGAGATGCGACCACAG |
| AN4663-PM-Seq-Rv | To sequence for checking the point mutation AN4663 E340A | TCGCCGGCATAATTCTTCAGTC |
| Ptef1-SeqInsert-F | To sequence for insertion using Ptef as promoter | CCTCGCTTCTCTCCTCCATC |
| Ttef1-SeqInsert-R | To sequence for insertion using Ttef as terminator | CAAGGTACACGCAAAACGAGGTAC |
| **PS name/number** | **Target** | **Sequence (5' à 3')** |
| **Target sequences involved in gene editing (Protospacers)** | | |
| PS328 | AN10909 | CGAGTGCATAATGTCCTTCA |
| PS329 | AN10909 | CCCACCACCTGCATTAGTTC |
| PS330 | AN2165 | AAATTTCCATGAGCAGATTC |

| | | |
|---|---|---|
| PS331 | AN2165 | CGCGAAGCAATGAACTGCTC |
| PS332 | AN1566 | TAAGGACAACTGGTCATCCG |
| PS333 | AN1566 | GGCTAGGGTATGTTAGGCTT |
| PS338 | AN0761 | ATCGGGAGGTGCTGAGATGA |
| PS339 | AN0761 | TCACCCTTCTGCCCTATTTG |
| PS340 | AN0134 | CAGTATGGAATCTGAAAGCG |
| PS341 | AN0134 | TACACTCAAGCATTAAGGAT |
| PS-AN4663-1 | AN4663 | GCTACCAAGAAGCTGGTAGT |
| PS-AN4663-2 | AN4663 | GATTATGAGGACTGTTTTGC |
| PS-AN3464-1 | AN3464 | ACGATGGCGTGGTACTGTTC |
| PS-AN3464-2 | AN3464 | TCAGTACCGCATTCGGCGGT |
| PS-AN2405-1 | AN2405 | CGCTCACAATGGCATCTGCA |
| PS-AN2405-2 | AN2405 | AGCGGATATAGCGGCACTAC |
| PS-AN7375-1 | AN7375 | AAGAAATCAGTCGAGCTCTC |
| PS-AN7375-2 | AN7375 | AACGGTGTTGCCTCTCAGTT |
| PS-AN10974-1 | AN10974 | TTGACATAGTCGACATTCGC |
| PS-AN10974-2 | AN10974 | TGAGTTTGGAAGCGGTTCCG |
| PS-AN9193-1 | AN9193 | TGCAGCGTCAGCCATGGTCA |
| PS-AN9193-2 | AN9193 | GCAGGTTCAATGTTGTGGTC |
| PS-AN8945-1 | AN8945 | TATTTTGTCGGTCGCAATGG |
| PS-AN8945-2 | AN8945 | GGCAGAAGCCGGAGTAGAGC |
| PS-AN4625-1 | AN4625 | GATTCAAGACACCTACTCCG |
| PS-AN4625-2 | AN4625 | CAGATCTATGCTGTCAAGGG |
| PS-AN6094-1 | AN6094 | CCTTCACAACAACCCATAAA |
| PS-AN6094-2 | AN6094 | CAGGTTGTACGTGATTTTCC |
| PS-AN9098-1 | AN9098 | ATCCAATTTACTCAGCTTCA |
| PS-AN9098-2 | AN9098 | GATATGGATATAACCGGGAT |
| PS-AN3096-1 | AN3096 | GCGGACATGGCGACTACTGC |
| PS-AN3096-2 | AN3096 | CGAAAGGGGAAGCCAGAACT |
| psAN2406-1 | AN2406 | AGGGCAGAAAACATCAATAT |
| psAN2406-2 | AN2406 | TATACAAGGTGCGAAGGTAG |
| PS-AN5630-1 | AN5630 | CATGGTCACCGAAAATGGCT |
| PS-AN5630-2 | AN5630 | CCACAATTTACTCACTCTTG |
| PS-AN5874-1 | AN5874 | CAACCTGACCAGATATTTGT |
| PS-AN5874-2 | AN5874 | GACCCGACAAGCCGAGATAT |
| PS-AN6713-1 | AN6713 | TTTGTAGAAATCGCGTGCCA |
| PS-AN6713-2 | AN6713 | TGAAGAGGAAACTGGAGGGG |
| PS-AN8849-1 | AN8849 | AAGAGAGCAACTAGCACGCA |
| PS-AN8849-2 | AN8849 | ATCAGATAGTGATTAATTTA |
| PS-AN10700-1 | AN10700 | TTCATCCACCGAGAAACTAA |
| PS-AN10700-2 | AN10700 | GAGGCTTTCAAGAAGTTTAG |
| PS-AN4663-point-mutation-2 | AN4663-E340 | GACCATCGTCGAAATCGACC |
| **Primers for *K. phaffii*** | | |
| 4817 | aA_PmeI_F | AAACGCTGTCTTGGAACCTA |
| 4818 | aA_PmeI_R | AAACTGTCAGTTTTGGGCCAT |
| 5552 | Fw_SapI_AN4663 | CGGGTTTGCTCTTCGCATACAACGACCGCACTTAATAATATCATCGCTG |
| 5359 | Rv_SapI_AN4663 | AAACCCGGGGCTCTTCGTTACCAGCCCTCCCAGACGCG |
| 5990 | FW-NativeSP_LsAA9A | ACGGCACTUTCCTTCGTCGCGTCAGCCGCGGCTCATACCCTCGTCTGGGGC |
| 5991 | RV-NativeSP_LsAA9A | AAGTGCCGUGAGCCCGAGGATGGAGTACTTCATCGTTTGGATCCTTCGAATAATTAGTTGTTTTTTGATC |
| 5701 | FW-USER-AN4663-episomal | ATTGAACAACUATTTCGAAACGATGGCGCCTTTTCGTTCAATTTATG |
| 5702 | RV-USER-AN4663-GAP-episomal | AGTTGTTCAAUTGATTGAAATAGGGACAAATAAATTAAATTTAAAGTC |

### Note S1 - DNA sequence LsAA9A

Native$^{SP}$-LsAA9A

```
        1 atgaagtact ccatcctcgg gctcacggca ctttccttcg tcgcgtcagc
cgcggctcat
       61 accctcgtct ggggcgtatg ggtcaacggc gttgatcaag gagatgggag
gaacatctac
      121 atcaggtctc ccccgaacaa taaccccgtc aagaacctga cttcgccgga
tatgacgtgc
      181 aatgtcgaca atcgcgttgt gcccaagagc gtgccagtca acgctggtga
tacgcttact
      241 ttcgaatggt accacaacac acgagacgac gatatcatcg catcttctca
tcacggaccc
      301 atcgccgtat acatcgcacc tgccgcttcg aacggtcaag ggaacgtttg
ggtgaagctg
      361 ttcgaggacg cctacaacgt caccaatagc acttgggctg tcgatcgtct
catcactgct
      421 catggacaac actccgttgt ggtcccccat gttgcacctg gagactacct
atttagagct
      481 gagatcattg cgctacatga ggcagattct ctgtactccc agaaccctat
ccgtggcgcg
      541 caattttata tctcgtgcgc tcaaatcacc atcaactctt ccgatgactc
gacacctctc
      601 cccgctggtg ttcccttccc cggtgcatac accgactcca cacccggtat
tcaattcaat
      661 atctacacta cccccgcgac aagctacgtt gctccccctc ccagcgtatg
gtctggagcg
      721 ttgggtgggt cgattgctca ggtcggagat gcttctctcg ag
//
```

### Note S2: DNA sequence TfAA10A

TfLPMO10A

```
        1 catgggtcgg tcatcaaccc cgcgacccgt aactacggtt gctggctgcg
ttggggccac
       61 gaccacctca accccaacat gcagtacgaa gaccccatgt gctggcaggc
ctggcaggac
      121 aaccccaacg ccatgtggaa ctggaacggc ctgtaccgcg actgggtcgg
cggcaaccac
      181 cgggctgccc tccccgacgg ccagctgtgc agcggtggcc tcaccgaagg
cggccgctac
      241 cgctccatgg acgccgtagg cccgtggaag accaccgacg tcaacaacac
cttcaccatc
      301 cacctgtacg accaggccag ccacggcgca gactacttcc tggtctacgt
caccaagcag
      361 ggcttcgacc cgaccaccca gccgctgacc tgggacagcc tggaactggt
gcaccagacc
      421 ggcagctacc ccccggccca gaacatccag ttcacggtcc acgcccccaa
ccgcagcggc
      481 cgccacgtgg tcttcaccat ctggaaggcc tcgcacatgg accagaccta
ctacctgtgc
      541 agcgacgtga acttcgtc
```

### Note S3: DNA sequence AN4663 in pBGP1

#### Codon variant AN4663

1 atggcgcctt ttcgttcaat ttatgagaaa gatgccacca aaaagcttgt tgtaggagca

61 gcgttgcttg tgctggctgc gttttatagt tatgtatttc tgctgactct ggcgcctgta

121 tatggttcta ctccttccca tatctttcac ggctatggag tcggtatcgc gggtgtagct

181 ggctggtttt cgaaggacat tgtggatcgc gtaagcggtc gtaaagcaat ctatgcaatt

241 ccagttttgg cgttcttttt accagtcgtt caatactttg tcagtcagca gtcatctgca

301 cttggcaatc ctgcggggcc aatcttcaca gaggtattgg ctctgtaccc tctggtcttg

361 ttatcagttg cttgtgcggg gaagctggtc caggccggtc ttaacttgca acgccatgga

421 gacttggtag ctgaacacat tcctttactt gggtcgtacg ttatttattc tgcaggggag

481 catcttatca aggccttctt atctcgcttc atcggctcca ccgtactttt gtcacgcgct

541 ggcttacaga ttcttatcgc aattttctac gctgccgccg ttccttccaa ggcgttactt

601 ctggccattc cagcgttctt attctcagtc acatcgaaca cccaccttcc tttgggccac

661 acaacgaccg cacttaataa tatcatcgct gatgacggct tcgcacttgt tgcacgccaa

721 gacagcacta ccgggtacat ttcagtcttg gataaccttg aggatgggtt tcgcgtaatg

781 cgctgtgacc acagtttact gggaggtcaa tggatcaaaa aacgtcccaa ttatactcct

841 ccagccgtaa aggaccccat ctatgcagta ttcacaatgt tggaggcagt gcgcttggtc

901 gaaacggctc acggtatccc ccgtgctgat gcgggctcca acgcgcttgt aatcggcttg

961 gggatcggca ctacccctgg cgctttgatt agtcacggaa ttgacaccac gattgtcgaa

1021 attgaccctg ttgtccacaa atatgccctt caatattttg accttcctga aaaccacaca

1081 cctatcattg aggacgcacg cgcttttgta cagcgttcgc gtaatgctcc acaacccaag

1141 cagtacgatt acattgtgca tgacgtgttc actggcggag ctgagccggt agagttgttc

1201 acctatgagt tcatctcggg cctgcatgcg cttttgaaag acgatggagt tatcgccatt

1261 aactacgcag ggatatttc cttatattca acagctctga gcatccgcac aatcaaaagc

1321 atttttccca cctgtcgctt gttccgcgag gctgccgccc cggagatcgg accggatttt

1381 acgaatatgg tcattttctg cacgaaatcg cgtggtgcac cgattacatt ccgcgatccc

1441 gtaccggaag atttcttggg aagccgcttt cgttctcgtt accttgttcc aaaacatgag

1501 gtagatgccg cgcaattcga caacgtcggg ttggaagacg gtcctcaggg acatggccgc

1561 cgcgtgctgg ttgacaaaga ggtcggtcgt ttacacaaat atcaggaccg ttccgcactg

1621 gagcattggg gaattatgcg taccgtcttg ccagatcgcg tctgggaggg ctgg

//
